# Systems analysis of long-term heat stress responses in the C_4_ grass *Setaria viridis*

**DOI:** 10.1101/2023.11.09.566437

**Authors:** Peng Zhang, Robert Sharwood, Adam Carroll, Gonzalo M Estavillo, Susanne von Caemmerer, Robert T. Furbank

## Abstract

A substantial number of C_4_ plants are utilized as food and fodder crops and often display improved resource use efficiency compared to C_3_ counterparts. However, their response to future extreme climates such as heatwaves is less understood. *Setaria viridis*, an emerging C_4_ model grass closely related to important C4 crops, was grown under high temperature for two weeks (42°C as compared to 28°C). High temperature resulted in stunted growth, but surprisingly had little impact on leaf area based photosynthetic rates. Rates of dark respiration significantly increased and there were major alterations in carbon and nitrogen metabolism in the heat-stressed plants, including reduced starch levels, accumulation of soluble sugars and an increase in leaf nitrogen content. Measurements of major phytohormones revealed a dramatic increase in abscisic acid in the heat-stressed plants. Leaf transcriptomics, proteomics and metabolomics analyses were carried out and mapped onto metabolic pathways of photosynthesis, respiration, carbon/nitrogen metabolism and hormone synthesis and signaling. Overall, upregulation of a number of stress-signaling pathways was observed, consistent with multiple potent signals leading to reduced plant growth. A systems model of plant response is presented based on oxidative stress, hormone and sugar signaling pathways.

## Introduction

C_4_ photosynthesis is a biochemical CO_2_ concentrating mechanism where CO_2_ is first assimilated by phosphoenolpyruvate carboxylase (PEPC) to produce a four-carbon compound, a C_4_ acid (Hatch and Slack, 1966; Hatch, 1987). This C_4_ acid diffuses into the bundle sheath (BS) cell where decarboxylation of the C_4_ acid releases the previously fixed CO_2_ in a cellular compartment surrounded by suberin that restricts leakage of CO_2_ (Danila et al., 2021). This creates a high local concentration of CO_2_ around Ribulose-1,5-bisphosphate carboxylase-oxygenase (Rubisco) so that the oxygenation reaction is suppressed, thus improving efficiency of CO_2_ assimilation while minimizing photorespiration. Most C_4_ plants have evolved the two-cell mechanism, where CO_2_ entering the mesophyll (M) cell is first converted to HCO ^-^ by carbonic anhydrase (CA), which is then assimilated into phosphoenolpyruvate (PEP) by PEPC to make oxaloacetate (OAA). Depending on the decarboxylation enzyme used, OAA is converted to malate or aspartate which is then transferred to adjacent BS cells where it is decarboxylated to release CO_2_ at the site of Rubisco (Hatch, 1987; Furbank, 2011).

C_4_ photosynthesis is considered an adaptation to environments that promote high photorespiration (Sage et al., 2018). The early lineages of C_4_ plants are thought to have evolved 25 – 30 million years ago, coinciding with the decline of atmospheric CO_2_ concentration from about 1000 ppm to well below present day levels (Sage, 2001; Zachos et al., 2001; Sage, 2004). In areas of the world where temperatures are frequently high, and drought and/or salinity are common, plants experienced high photorespiratory flux and were under selective pressure to evolve mechanisms that allow more efficient carbon gain (Sage et al., 2018). Presently, over 60 independent lineages of C_4_ plants have been discovered, and altogether contributing towards ∼25% of the terrestrial primary productivity (Edwards et al., 2010). Some of the economically important C_4_ crops include maize, sorghum, millet and sugar cane. Given that C_4_ plants are regarded as relatively tolerant of higher growth temperatures (Berry and Bjorkman, 1980), it is not surprising that the majority of studies on longer term effects of high temperature stress on photosynthesis, metabolism, growth and yield have been carried out on C_3_ plants (Sgobba et al., 2015; Kask et al., 2016; Hütsch et al., 2019; Haider et al., 2021; He et al., 2022).

Plant heat stress response during vegetative growth is extremely complex. The rates of most enzymatic reactions are sensitive to temperature increases, and heat stress can directly impact on protein and membrane stability, causing changes in the rates of biological processes. Heat may also cause the production of reactive oxygen species (ROS), which may act as signals to halt/alter plant growth but may also cause oxidative damage to the plant tissues leading to senescence and death (Bhattacharjee, 2005; Sharkey, 2005; Suzuki and Mittler, 2006). Heat stress can significantly decrease plant photosynthetic capacity by negatively impacting on both the light and dark reactions. The structure and function of photosystem (PS) II is particularly heat labile (Sharkey, 2005). Severe heat stress (above 45 °C) has been shown to directly cause structural damage to PS II, including dissociation of the oxygen evolving complex (Papageorgiou and Murata, 1995), and dissociation of the reaction centre protein complex (Nash et al., 1985). Moderate heat stress did not appear to severely affect PS II structure but was shown to inhibit the repair of photodamaged PS II by inhibiting the *de novo* synthesis of D1 protein, causing disruption to linear electron transport (Takahashi et al., 2004; Sharkey, 2005; Allakhverdiev et al., 2008b; Allakhverdiev et al., 2008a).

At elevated temperatures, both C_3_ and C_4_ Rubisco has been shown to be subject to deactivation due to heat labile nature of Rubisco activase (RCA) and the build-up of inhibitors that reduce Rubisco activity (Sharkey et al., 2001; Crafts-Brandner and Salvucci, 2002; Salvucci and Crafts-Brandner, 2004). In addition, the thermal lability of other Calvin-Benson cycle enzymes may also contribute to reduced carbon fixation under heat (Berry and Bjorkman, 1980; J Berry and Bjorkman, 1980; Allakhverdiev et al., 2008a). Heat stress has been shown to inhibit photosynthesis in C_4_ crop species such as sugarcane (40 °C), maize (above 38 °C) and sorghum (40 °C) with carbon fixation appearing to be more sensitive than light harvesting and electron transport (Blum, 1986; Crafts-Brandner and Salvucci, 2002; Wahid, 2007; Perdomo et al., 2017). Furthermore, short-term heat stress (4 hours) at 40°C has been shown to inhibit photosynthesis in *S. viridis* plants (Anderson et al., 2021).

In addition to the direct impact of heat on photosynthetic components, high temperature may affect carbon fixation by influencing stomatal conductance. Under prolonged heat stress, plants may initially up-regulate transpiration for evaporative cooling of the leaves, particularly if vapor pressure deficit is high. If availability of soil water is limited, this increase in transpirational water loss may induce water stress in the heat-stressed plants. Under such circumstances, a range of drought-related responses will be induced including stomata closure, which can limit CO_2_ availability to the chloroplast and inhibit photosynthesis. Water stress has been frequently observed in plants experiencing high temperature (Tsukaguchi et al., 2003; Wahid and Close, 2007).

The impact of heat stress on plant growth depends on the plant species and the developmental stage. During vegetative growth, heat stress may inhibit cell expansion by lowering the water potential, leading to reduction in cell and overall plant size (Bañon et al., 2004; Potters et al., 2007). The developmental pattern may also be changed by heat stress. Changes in stomata and trichome densities and number of xylem vessels in both shoots and roots have been observed in *Lotus* plants (Bañon et al., 2004). Heat stress may inhibit the expansion of the first internode, which was observed in sugarcane plants grown under high temperature (Ebrahim et al., 1998). These plants also exhibited increased tillering, early senescence and reduced biomass accumulation (Ebrahim et al., 1998). The impact of high temperature on enzymes directly involved in plant cell wall expansion is not well understood. In many cases, the reduction in plant growth caused by heat is thought to be the result of photosynthesis inhibition and altered assimilate partitioning (Wahid et al., 2007).

Given the advancement of various ‘omics technologies, and the huge complexity of plant heat stress response, there is a clear need for experiments employing a systems approach, by integrating multiple omics platforms to gain a holistic understanding of plant heat stress response and tolerance mechanisms in plants carrying out C_4_ photosynthesis. An experimental system was established to explore the long-term heat stress response of *S. viridis* seedlings. Seedlings were initially established under normal growth conditions (28 °C day / 20 °C night) for two weeks, then subjected to a heat stress treatment by transferring to a high temperature growth chamber (42 °C day / 32 °C night) for another two weeks until the completion of vegetative growth and the onset of flowering. We consider this heat stress treatment “long-term” because the length of the treatment corresponds to approximately half of the vegetative growth phase. Physiological, biochemical, transcriptomics, proteomics and metabolomics data were measured in control and heat-treated plants and compared to determine mechanisms of perception of the stress and the plant system response.

## Materials and Methods

### Plant growth and anatomical measurements

Seeds of *Setaria viridis* genotype A10 were treated in 10% liquid smoke for 2 hours at room temperature to promote germination. Seeds were sown onto a rice mix (prepared by CSIRO) in a small clear plastic container (10cm x 10 cm x 5 cm) with the lid closed to prevent drying. Seeds were germinated in a Conviron growth chamber under 16h/8h day-night cycle, and the temperature was set at 28 °C day/20 °C night, relative humidity 40-60%, CO_2_ at 380 ppm, and an irradiance at 1000 μmol quanta m^-2^ s^-1^. After germination, seedlings were transplanted into individual pots made with PVC plastic pipe that are 10 cm in diameter and 75 cm tall containing the same rice mix. At two weeks old, 20 plants were transferred into a Conviron growth chamber that had the exact same condition except the temperature was set at 42 °C day/32 °C night. Plants were watered daily in the morning and fertilized with diluted Seasol^TM^ liquid fertilizer 3 times a week.

At the end of the treatment, three plants from the control and heat-stressed group were taken for destructive analysis. The roots were washed and the roots and shoots cut separately with scissors and placed in paper bags. The material was dried in an 80 °C oven for 3 days and the weight of the dried material measured. One fully expanded fresh leaf from each plant was taken for fixation for anatomical analysis. The leaf was equally divided into tip, middle, and bottom sections, and placed in 25% glyceraldehyde solution in glass vials. Leaf sections were finely sliced by hand and visualized using a microscope attached to a camera.

### Gas exchange measurements

Gas exchange was measured on the flag leaf of the 4-week-old control and heat-stressed plants using a LI-6400XT with a LED light source (Licor). Measurements were performed early in the morning, from ∼2 hours into the day cycle. Measured leaves were initially equilibrated under standard condition of 380 μmol.mol^-1^ CO_2_, 25 °C leaf temperature, 2000 μmol quanta m^-2^. s^-1^, and a flow rate of 500 μmol.s^-1^, until leaf photosynthesis stabilized. Photosynthetic CO_2_ response (A-C_i_) curves were performed by first step wise decrease in CO_2_ partial pressure from 400 μmol mol^-1^ to 0 μmol.mol^-1^, then step wise increase from 0 μmol.mol^-1^ to 1500 μmol.mol^-1^, while maintaining leaf temperature at 25 °C and irradiance of 2000 μmol.m^-2^.s^-1^. Photosynthetic light response curves were performed by step wise increasing the light intensity from 0 μmol quanta m^-2^.s^-1^ to 2000 μmol.m^-2^. s^-1^, while maintaining CO_2_ partial pressure at 380 μmol.mol^-1^ and leaf temperature at 25 °C. Leaf dark respiration measurements were performed with dark adapted plants that just exited the night cycle. Leaves were first measured at 25 °C, 380 μmol.mol^-1^ CO_2_. Then leaves were measured at 20, 25, 30, 35, 40, 45 °C. Measurements were taken at each temperature when leaf dark respiration stabilized in the new condition. At the end of the gas exchange experiments, leaf disks were taken from the measured leaves (4 plants for each condition) and snap-frozen in liquid nitrogen for measurement of enzyme activity, chlorophyll content, starch content, and protein content.

### Enzyme activity assays

Rubisco and PEPC assays were performed as previously published (Sharwood et al., 2016). Leaf discs (0.5cm^2^) were punched from the mid-section of fully expanded leaves from the control and heat stressed plants were collected and snap-frozen in liquid nitrogen. One leaf disc was homoghenised with a glass homogeniser in 500 μL of freshly prepared ice-cold extraction buffer (100 mM HEPES-KOH, pH 7.4, 5 mM DTT, 0.1% BSA, 0.05% Triton X-100, 2 mM EDTA, 5 mM MgCl_2_, 1% PVPP) plus 10 μL of protease inhibitor cocktail (P-9599; Sigma, St. Louis, MO, USA). The homogenate was quickly transferred into a 1.5 mL tube and centrifuged at top speed for 30 seconds at 4 °C. To cuvette 1 and 2, 435 μL of freshly prepared Rubisco assay buffer (100 mM EPPS, pH 8, 20 mM MgCl_2_, 1 mM EDTA, 10 mM NADH, 10 mM ATP, 50 mM coupling enzyme, 400 mM NaHCO3) was added; to cuvette 3, 435 μL of freshly prepared PEPC assay buffer (100 mM EPPS, pH 8, 20 mM MgCl2, 1 mM EDTA, 40 mM NADH, 5 mM glucose-6-phosphate, 400 mM NaHCO3, 0.5 unit of malate dehydrogenase enzyme) was added. Enzyme activity was determined by the consumption of NADH at 340nm using a diode array spectrophotometer at 25°C (Agilent).

### Leaf elemental analysis and chlorophyll and carbohydrate content

Leaf chlorophyll content of the leaves was measured following the method developed by Porra et al. (1989). Leaf tissue was homogenised using a glass homogeniser in ice-cold 80% acetone buffered with 2.5 mM phosphate buffer (pH 7.8). The homogenate was centrifuged at 12,000 x g for 10 min at 4 °C, and 100 μL of the supernatant was diluted with 900 μL of the extraction buffer before reading the absorbance at 750 nm, 663.6 nm and 646.6 nm on a spectrophotometer.

Leaf discs from the heat stress experiment were oven-dried and wrapped into a small aluminium weighing boat with their accurate weight recorded. The percentage of carbon and nitrogen relative to leaf dry weight was estimated by combustion of samples in an elemental analyser (EA1110, Carlo Erba).

### RNA extraction and quality analysis

Total RNA was extracted from frozen leaf powder (from pooled mid-section of fully expanded leaves) using TRIzol™ Reagent. For each tube of homogenized leaf sample (∼100 mg), 1 mL of TRIzol™ Reagent was mixed in and incubated at room temperature for 5 min to lyse the cells. The cell lysate was centrifuged at 12,000 x g for 15 min at 4 °C to remove debris, and the supernatant was transferred to a new tube. To each tube 0.2 mL of chloroform was added, and the tubes were shaken by hand for 15 seconds and incubated for 2-3 min at room temperature. The mixture was centrifuged at 12,000 x g for 15 min at 4 °C, and the upper aqueous phase containing the RNA was transferred into a new tube. For RNA precipitation, 0.5 mL of isopropanol was added to the aqueous phase and incubated at room temperature for 10 min. The mixture was centrifuged at 12,000 x g for 10 min at 4 °C and the supernatant was discarded. The resulting RNA pellet was washed with 75% ethanol and centrifuged at 7,500 x g for 5 min at 4 °C. The wash was repeated three times. The RNA pellet was resuspended in 50 μL of RNAse-free water.

The quality and accurate concentration of the extracted RNA was measured by Agilent 2100 Bioanalyser using the Agilent RNA 6000 Nano Kit following the manufacture’s instruction.

### RNA Sequencing and Library Preparation

Isolated RNA sample was sent to the Next Generation Sequencing Facility at the Hawkesbury Institute for the Environment at the Western Sydney University for RNA Seq library construction and sequencing. Ribosomal RNA was removed before the construction of strand-specific RNA library using Illumina TruSeq RNA Sample Preparation kit version 2. Data was obtained for five control samples and six heat-stressed samples.

### RNA Sequencing Data analysis

The quality of the RNA Seq data was checked using the FASTQC program (https://www.bioinformatics.babraham.ac.uk/projects/fastqc/). Reads were quality trimmed and adapter sequences were removed using Trimmomatic (Bolger et al., 2014) with the following setting: LEADING:10 TRAILING:10 SLIDINGWINDOW:4:15 MINLEN:50 (and the rest of the settings are default). This setting means the first and the last 10 bases are cut off if their quality score is below the threshold; the average score of a sliding window size of 4 bases is calculated and the read trimmed if the average score falls below 15; read is discarded if read length is below 50. The quality trimmed reads were mapped to the genome of *Setaria italica* (v2.1) (downloaded from: http://plants.ensembl.org/) using TopHat2 with the default setting allowing 2 mismatches (Kim et al., 2013). The alignments were counted to exons using HT-Seq with mode set to union (Anders et al., 2015). The read count data was analysed in DESEQ for differential expression analysis (Anders and Huber, 2010). Differential expression analysis was performed and fold-changes (heat-stress over control) was calculated and log2 transformed. Multiple testing correction was performed by the Benjamini-Hochberg procedure and FDR set to 5%. Annotation file for *Setaria italica* genome was downloaded from Phytozome (https://phytozome.jgi.doe.gov/pz/portal.html#). The file contains the orthologous gene information for each *Setaria* gene, orthologous gene from *Arabidopsis*, *Rice* and *Maize* was assigned to each *Setaria* gene. GO terms were assigned to each gene using the rice GO annotation, and the genes were further annotated with Mapman mappings (http://mapman.gabipd.org/mapmanstore). The raw data and metadata is uploaded to GEO, Accession Number: GSE216993. Processed data is presented in Supplementary Data File 1.

### Proteomics analysis

Total leaf protein was extracted using the TCA/acetone method (Damerval et al., 1986). Homogenized leaf tissue (∼300 mg from each sample) from the heat stress experiment (same sample used for RNA extraction) was extracted with 1 mL of cold precipitation buffer (10% w/v trichloroacetic acid, 0.07% v/v 2-mercaptoethanol in acetone, pre-chilled at -20 °C overnight). The mixture was incubated at -20 °C for at least 2 hours (or overnight). The homogenate was centrifuged at 19,000 x g for 15 min at 4 °C and the supernatant was discarded using a pipette. The pellet was washed three times with ice-cold acetone (with 0.07% 2-mercaptoethanol) to remove chlorophyll, and the pellet was dried under nitrogen gas. The pellet containing protein was resuspended in solubilization buffer (8 M urea, 2% w/v CHAPS, 30 mM Tris-HCl pH 7.8, use approximately 60 μL per milligram of protein) with the help of a sonicator water bath. The mixture was centrifuged at 14,000 x g to remove insoluble debris, and the supernatant containing protein was collected. The extracted protein solution was quantified using the 2-D Quant Kit (GE Healthcare) following the manufacturer’s instruction.

Disulfide bonds were reduced by adding solubilisation buffer to 2mg of protein to a total volume of 995 μL. Fresh DTT (1 M) was made by dissolving a small amount of powder in 50 mM ammonium bicarbonate (NH_4_CO_3_), and 5 μL of 1 M DTT was added to each tube of protein solution. The solution was incubated at 37 °C for 1 hour for complete reduction. During the incubation iodoacetamide to 40mM was added (Pieterse et al., 2000) and incubated in the dark for 2 hours at room temperature. An additional 5 μL of 1 M DTT was added to the tubes to stop the alkylation, the tubes were incubated at room temperature for 15min. For protein digestion, the proteinase enzyme Lys-C (Pierce™ Lys-C, MS grade) was added at a 1 : 100 enzyme : protein (w/w) ratio, and incubated at 37°C for 4 hours. The samples were diluted 1:8 with 50 mM NH_4_CO_3_ to reduce the urea concentration to 1 M. CaCl_2_ was also added to each sample to a final concentration of 1 mM. The proteinase enzyme Trypsin (Pierce™ Trypsin, MS grade) was added to the tubes at an enzyme : protein ratio (w/w) of 1 : 30, and incubated in a 37 °C water bath overnight. The samples were cooled on ice, and the digestion was stopped by adding trifluoroacetic acid (TFA) to 1% v/v. The tubes were centrifuged at 2,500 x g for 10 min at room temperature, and the supernatant was collected. The volume was reduced by drying the samples in a SpeedyVac and stored at -20 °C for further processing.

Dried peptide samples were reconstituted in 1 mL of 5% formic acid. The reconstituted peptides were labelled following the on-column stable isotope dimethyl labelling protocol (Boersema et al., 2009).

The labeling reagent was prepared by combining 4.5 mL of sodium phosphate buffer pH 7.5 (prepared by mixing 1 ml of 50 mM NaH_2_PO_4_ with 3.5 mL of 50 mM Na_2_HPO_4_) with 250 μL of 4% (v/v) formaldehyde in water (CH_2_O or CD_2_O) and 250 μL of 0.6 M sodium cyanoborohydride in water (NaBH_2_CN or NaBD_3_CN). For the heat-stress experiment the light label was introduced to the peptide samples from the control plants, and the intermediate label was introduced to the peptide samples from the heat-stressed plants. Sep-Pak C18 cartridges (Waters) were engaged onto a vacuum extraction manifold (Waters). Each cartridge was washed with 2 mL of acetonitrile and conditioned with 2 mL of reverse-phase solvent A (0.6% (v/v) acetic acid). Peptide samples (500 μg equiv.) were loaded onto each cartridge, and the cartridge was washed with 2 mL of reverse-phase solvent A. The labelling reagent was added to the cartridges, 1 mL at a time for a total of 5 mL for each sample and was allowed to flow through the cartridges slowly. The cartridges were again washed with 2 mL of reverse-phase solvent A, and the labelled peptide sample was eluted with 500 μL of reverse-phase solvent B (0.6% (v/v) acetic acid and 80% (v/v) acetonitrile) and collected into a Lo-bind protein tube.

### Mass spectrometry

The peptide samples were dried in a SpeedVac and submitted to the Sydney Mass Spectrometry facility (The Charles Perkins Centre, The University of Sydney) for untargeted mass spectrometry analysis. Reconstituted peptide (1 μL) was injected into an in-house built fritless nano 75 μm × 30 cm column packed with ReproSil Pur 120 C18 stationary phase (1.9 μm, Dr, Maisch GmbH, Germany) fitted on a Thermo Scientific Ultimate 3000 HPLC and auto-sampler system for separation, before introduced into a Q Exactive Plus mass spectrometer (Thermo Fisher Scientific, Waltham, MA, USA). Peptide fractions were directed for MS through a nano-electrospray interface, and charged species were captured and the Orbitrap mass analyser was used to acquire full MS with m/z range of 350-1550. The 20 highest peaks were fragmented by applying C-trap dissociation collision and tandem MS were generated. The mass spectrometry acquisition time was 60 min in total for each sample.

### Proteomics data analysis

The generated “.raw” files containing acquired data was analysed using the freely available proteomics software MaxQuant (version 1.5.2.8) (Cox and Mann, 2008; Tyanova et al., 2016a). The *Setaria italica* protein sequence v2.1 (Bennetzen et al.,2012) (downloaded from: http://plants.ensembl.org/) was used for library searching of the peptides. Peptide modifications selected include DiMethy Lys0/ DiMethy N-term0 (for light labelled peptides), DiMethy Lys4/ DiMethy N-term4 (for intermediate labelled peptides), and the level of multiplicity is 2, to achieve quantitation. The mass error tolerance for the main peptide search was 10 ppm. Other search parameters are kept default as recommended in Tyanova et. al. 2016. About 1800 protein groups with quantitative information (the ratio between intermediate and light labelled peptides) was obtained and further analysed in the Perseus software (Tyanova et al., 2016b). Mainly, the protein ratio of log2 transformed, statistical testing was performed using Student’s T-test with a FDR of 0.1. Additionally, when a protein group was only detected in less than 3 out of the 5 samples submitted, the ratio was determined as insignificant and not used. Processed data is presented in Supplementary Data File 1 and raw data uploaded to PRIDE.

### Metabolite Analysis

About 100 mg of frozen leaf powder samples (the same material used for RNA and protein analysis) for extraction was weighed into a pre-frozen 2 mL Eppendorf tube containing a steel ball bearing without thawing. An accurate weight was recorded for each sample for use later in data normalization. The tissues were further homogenized in a TissueLyser II bead mill (Qiagen; Cat. No. 85300) for 1 min at 20 Hz without thawing out. Then, to each tube containing frozen tissue powder was added 5 volumes (5 μL per mg fresh weight of tissue) of cold (-20°C) Metabolite Extraction Medium (85% (v/v) HPLC grade MeOH (Sigma), 15% (v/v) untreated MilliQ H_2_O, 8.7 µg mL^-1^ ribitol as internal standard), tubes were vortexed briefly, rapidly transferred to an Eppendorf Thermomixer Comfort (Eppendorf; Cat. No. 5355 000.011) and heated to 65°C and shaken at 1400 RPM for 15 min. Tubes were then centrifuged at 20,000 x g for 10 min to pellet insoluble material and 80% of the supernatant was transferred into a new clear 2 mL round-bottom polypropylene Eppendorf tube and stored as a stock extract at -80°C until further analysis.

From each metabolite extract, 15 μL was aliquoted into a glass vial insert and dried in a centrifugal vacuum concentrator (Labconco CentriVap, Catalog Number 7811030). Dried metabolite extracts were chemically derivatized by methoximation and trimethylsilylation on a Gerstel MPS2XL Multipurpose Sampler (Gerstel) operating in the PrepAhead mode for automated online derivatization and injection. The derivatization procedure consisted of the following steps: (1) addition of 10 μL of 20 mg ml^−1^ methoxyamine hydrochloride (Supelco, Catalog Number 33045-U) in anhydrous derivatization-grade pyridine (Sigma-Aldrich, Catalog Number 270970) and incubation at 37°C for 90 min with agitation at 750 RPM; (2) addition of 15 μL of derivatization grade N-methyl-N-(trimethylsilyl)trifluoroacetamide (MSTFA; Sigma-Aldrich; Catalog Number 394866) and incubation at 37°C for 30 min with agitation at 750 RPM; 3) addition of 5 μL of alkane mix [0.006% (v/v) n-dodecane, 0.006% (v/v) n-pentadecane, 0.006% (w/v) n-nonadecane, 0.006% (w/v) n-docosane, 0.006% (w/v) n-octacosane, 0.006% (w/v) n-dotriacontane, and 0.006% (w/v) n-hexatriacontane dissolved in anhydrous pyridine and incubation for 1 min at 37°C with agitation at 750 RPM. Samples were injected into the GC/MS instrument immediately after derivatization.

Derivatized metabolite samples were analysed on an Agilent 5975C GC/MSD system comprised of an Agilent GC 7890N gas chromatograph (Agilent Technologies, Palo Alto, CA, USA) and 5975C Inert MSD quadrupole MS detector (Agilent Technologies, Palo Alto, CA, USA). The GC was fitted with a 0.25 mm ID, 0.25 μm film thickness, 30 m Varian FactorFour VF-5 ms capillary column with a 10 m integrated guard column (Varian, Inc., Palo Alto, CA, USA; Product No. CP9013). Samples were injected into the split/splitless injector operating in splitless mode with an injection volume of 1 μL, an initial septum purge flow of 3 mL min^−1^ increasing to 20 mL min^−1^ after 1 min and a constant inlet temperature of 230°C. Helium carrier gas flow rate was held constant at 1 mL min^−1^. The GC column oven was held at the initial temperature of 70°C for 1 min before being increased to 325°C at 15°C min^−1^ before being held at 325°C for 3 min. Total run time was 21 min. Transfer line temperature was 250°C. MS source temperature was 250°C. Quadrupole temperature was 150°C. Electron Impact ionization energy was 70 eV and the MS detector was operated in full scan mode in the range of 40– 600 m/z with a scan rate of 3.6 Hz. The MSD was pre-tuned against perfluorotributylamine (PFTBA) mass calibrant using the “atune.u” autotune method provided with Agilent GC/MSD Productivity ChemStation Software (Revision E.02.01.1177; Agilent Technologies, Palo Alto, CA, USA; Product No. G1701EA).

### GC/MS data processing

All GC/MS data were processed using the online MetabolomeExpress data processing pipeline4 (Carroll et al., 2010). Raw GC/MS files were exported to NetCDF format using Agilent MSD ChemStation software (Revision E.02.01.1177; Agilent Technologies, Palo Alto, CA, USA; Product No. G1701EA). Peak detection settings were: Slope threshold = 200; Min Peak Area = 1000; Min. Peak Height = 500; Min. Peak Purity Factor = 2; Min. Peak Width (Scans) = 5; Extract Peaks = on. Peaks were identified by MSRI library matching which used retention index and mass-spectral similarity as identification criteria. MSRI library matching parameters were as follows: RI Window = ± 2 RI Units; MST Centroid Distance = ± 1 RI Unit; Min. Peak Area (for peak import): 5000; MS Qualifier Ion Ratio Error Tolerance = 30%; Min. Number of Correct Ratio Qualifier Ions = 2; Max. Average MS Ratio Error = 70%; Remove qualifier ion not time-correlated with quantifier ion = OFF; Primary MSRI Library = “Carroll_2014_Arabidopsis_Photorespiration_Mutants.MSRI”; Add Unidentified Peaks to Custom MSRI Library = OFF; Use RI calibration file specified in metadata file = ON; Carry out per-sample fine RI calibration using internal RI standards = OFF.

The Carroll_2014_Arabidopsis_Photorespiration_Mutants.MSRI primary library contains entries derived manually from analyses of authentic metabolite standards run under the same GC/MS conditions as the biological samples as well as entries for unidentified peaks that were automatically generated by MetabolomeExpress while processing the data from the reference photorespiration mutants. Library matching results were then used to construct a metabolite x sample data matrix with peak areas being normalized to internal standard (i.e., ribitol). As a quality control filter, samples were checked for the presence of a strong ribitol peak with a peak area of at least 1 × 10^5^ and a deviation from the median internal standard peak area (for that GC/MS batch sequence) of less than 70% of the median value. Statistical normalization to tissue mass was not required because chemical normalization to tissue mass had already been carried out by adjusting extraction solvent volume proportionally to tissue mass. The heat Vs. control signal intensity ratio of each metabolite was calculated by dividing the mean (normalized) signal intensity of each metabolite in the heat-stressed plants by its mean (normalized) signal intensity in the control plants. Statistical significances were calculated by two-tailed Welch’s t-tests (n = 5) in the MetabolomeExpress Comparative Statistics tool. The full dataset has been uploaded to the MetaboLights metabolomics data repository (Haug et al., 2020) under the accession number: MTBLS8842. Processed data is presented in supplementary data file 2.

### Hormone analysis using UPLC-Orbitrap

Samples from five biological replicates for the control and five biological replicates for the heat-stressed plants were analysed. The frozen tissue samples (∼ 100 mg weighed into a tube with fresh weight recorded) were homogenized in a TissueLyser II bead mill (Qiagen; Cat. No. 85300) for 1 min at 20 Hz without thawing out. For auxin extraction (Ng et al., 2015), 20 μL of the internal standard (1 μg/mL of 3-[2H5]indolylacetic acid) followed by 1 mL extraction solvent (20% methanol:79% propanol:1% glacial acetic acid) were added to each tube, and the tubes were incubated in a sonicator bath for 15 min at 4°C. Samples were then centrifuged at 16,100 x g for 15 min. The supernatant was transferred to a fresh tube and dried in a centrifugal vacuum concentrator (Labconco Centrivap). Extraction was repeated and the supernatant combined with the first batch and dried again. Dried samples were resuspended in methanol/water (60:40, v/v) and filtered with a Nanosep MF GHP (hydrophilic polypropylene) 0.45-μm filter (Pall Life Sciences) prior to injection. For extraction of ABA, JA and SA (Miyazaki et al., 2014; Xu et al., 2016), 20 μL of the internal standards ([^2^H_6_]ABA or 1 μg mL^-1^ dihydrojasmonic acid and 2-hydroxybenzoic acid) followed by 950 μL of extraction solvent (70 : 30, acetone : 50 mM citric acid) was added to each tube of frozen leaf powder. Tubes were placed on a shaker at 4°C in the dark for 5 h, and then the tubes were left uncapped in a fume hood to allow the acetone layer to evaporate overnight. The ABA, JA and SA in the remaining aqueous phase were extracted by partitioning three times with 500 μL of diethyl ether. The ether phase was collected in a 2 mL glass vial (Kinesis Australia Pty, Redland Bay, Qld) and evaporated using a centrifugal vacuum concentrator (Labconco Centrivap) until dry. Dried samples were resuspended in 60% methanol (50 μL) and filtered with a Nanosep MF GHP (hydrophilic polypropylene) 0.45-μm filter (Pall Life Sciences) prior to injection. Samples and standards (5 μL) were injected onto an Agilent Zorbax Eclipse 1.8 μm XDB-C18 2.1 × 50 mm column. For ABA, SA and JA, the column temperature was held constant at 45 ± 0.5 °C. Solvent A consisted of 0.1% aqueous formic acid, and solvent B consisted of methanol with 0.1% formic acid. JA, SA and ABA were eluted with a linear gradient from 10% to 50% solvent B over 8 min, 50% to 70% solvent B from 8 min to 12 min (then held at 70% from 12 min to 20 min) at a flow rate of 200 μL/min. For auxins, the column temperature was 35 ± 0.5 °C and the linear gradient was the same as above using solvent A consisting of 0.1% aqueous formic acid and solvent B of 90% methanol/water with 0.1% formic acid. Solvents were LC-MS grade from Fisher Chemical. The eluted phytohormones from the column were introduced into the mass spectrometer via a heated electrospray ionsiation (HESI-II) probe and analysed using the Q-Exactive Plus Orbitrap (Thermo Scientific, Waltham, MA, USA). The HESI was operated in the negative ion polarity for JA, SA and ABA with parameters as follow: the electrospray voltage was 2.5kV, and the ion transfer tube temperature was 250 °C. The vaporized temperature and the S-lens RF level were 300 °C and 50 V, respectively. Ultra-high purity nitrogen as used as the sheath gas, auxiliary gas and sweep gas at flows of 45 L/min, 10 L/min and 2 L/min, respectively. For auxins, the HESI was operated in the positive ion polarity and the electrospray voltage was set to 3.5kV. Positive or negative ion polarity tandem mass spectrometry was carried out using targeted parallel reaction monitoring (PRM) with a mass resolution of 17,500 at 1.0 microscan. The AGC (Automatic Gain Control) target value was set at 1.0E+05 counts, maximum accumulation time was 50 ms and the isolation window was set at m/z 4.0. Data were acquired and analysed using the Thermo Scientific Xcalibur 4.0 software.

### Gene Ontology (GO) enrichment analysis

The list of gene IDs for significantly up- and down-regulated genes (padj < 0.05) in the heat-stressed plants were submitted to the agriGO website (bioinfo.cau.edu.cn/agriGO) for functional characterization using its singular enrichment analysis function. The Setaria italica v2.1 database was used to assign GO terms to genes. Only the list of expressed genes (mean RPKM > 20 in both of the conditions) was used as the reference background when considering enrichment. A GO category was considered significantly over-represented when the hypergeometric test p < 0.05, and a Benjamini-Yekutieli FDR < 0.1, and at least five entries mapped. To reduce the complexity of the GO dataset, the list of significant GO terms in the Biological process category was input into REViGO (Supek et al., 2011) to remove functional and semantic redundancies. Hierarchical clustering was further performed with the refined list of GO in REViGO using the default setting.

### Global analysis of hormone-regulated genes

The transcriptional responses of hormone-regulated genes in the heat stressed Setaria plants were compared to the hormone-associated responses in Arabidopsis. The Arabidopsis hormone treatment datasets were kindly curated and provided by Dr Michael Groszmann, which he obtained from the AtGenExpress database (atpbsmd.yokohama-cu.ac.jp/AtGenExpressJPN/AtGenExpress.html). This data consists of microarray-based gene expression responses to exogenous treatment by various hormones in Arabidopsis seedlings. The method of analysis was adopted from Groszmann et al. (2015) with variations to address species differences. To perform the comparison between Setaria and Arabidopsis, the list of expressed (RPKM >20) Setaria gene IDs were converted to Arabidopsis gene IDs by identifying the closest related orthologue. This was done using the *S. italica* genome annotation file version 2.1 from Phytozome (https://phytozome.jgi.doe.gov/pz/portal.html). The list of orthologous Arabidopsis gene IDs was mapped to the list of expressed genes from the Arabidopsis hormone treatment microarray dataset to identify a common list of genes expressed in both experiment (Arabidopsis hormone and Setaria heat), forming the “working set”. For analysis of each hormone response, the list of up-regulated (stimulated) and downregulated (repressed) genes in response to a particular hormone treatment was analysed separately. Count data was obtained for hormone-stimulated genes that were also up-regulated in the heat experiment, and for hormone-stimulated genes that were down-regulated experiment, under the condition that the genes exist in the “working set”, so that only genes expressed in both experimental conditions were compared. Similarly, count data was obtained for hormone-repressed genes by comparing to up-regulated and down-regulated gene lists from the heat experiment. To avoid the situation where multiple Setaria genes can be mapped to the same Arabidopsis ID, thereby inflating the count due to gene duplication, redundancy in Arabidopsis ID in the up- and down-regulated gene list from the heat experiment was removed, so that each Arabidopsis gene was only compared once in each scenario. The count data formed the basis of contingency tables for carrying out Fisher’s exact test, to identify overrepresentation of hormone-regulated genes in the Setaria heat stress treatment (p<0.01 was considered significant).

## Results

### Response of growth and photosynthesis to high temperature

Growth of *S. viridis* at 42°C/32°C day/night cycle resulted in a marked stunting of plant growth (Figure 1A). Both the shoot and the root dry biomass were reduced by approximately 50% compared to the plants grown at 28°C/22°C, while the root to shoot ratio was largely unaffected (Figure 1B,C).

**Figure 1.**
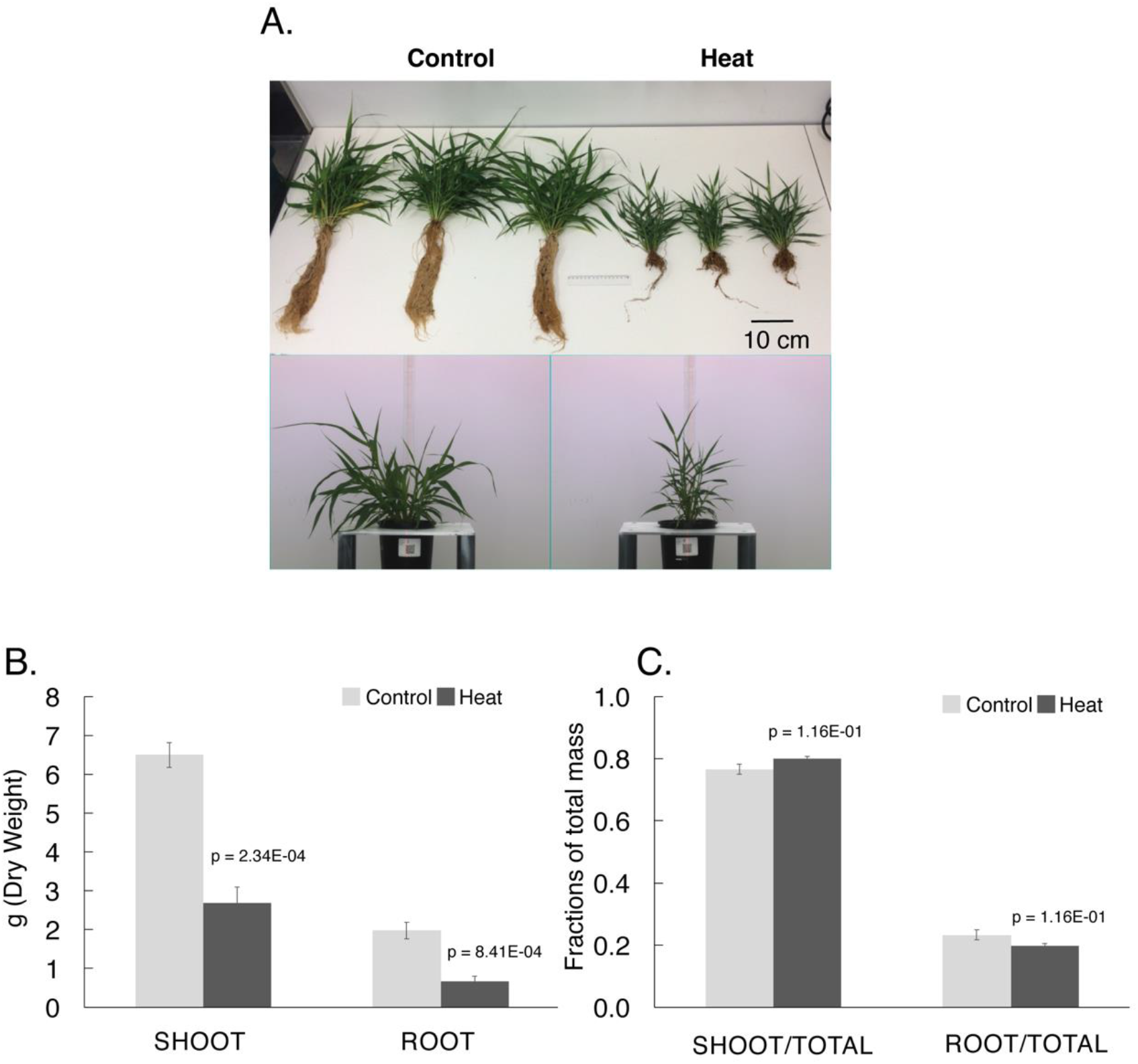
The growth and biomass accumulation of S. viridis in response to heat stress. A) Representative photos of plants grown at 28 °C day/22 °C night (Control) and 42 °C day/32 °C night (Thoreen et al., 2012). B) The shoot and root dry biomass, and (C) Biomass allocation to the shoot or root as a fraction of total biomass in the control and heat-stressed plants. Error bars shown represent the standard error of the mean (SEM), and p-values were calculated from Student’s t-tests (N = 3).

Anatomical studies showed the youngest fully expanded leaves were smaller in heat stressed plants, but leaf thickness was not significantly different, while interveinal distance was slightly reduced (Supplementary Figure 1).

Despite the significant effects on plant growth and development, growth at high temperature had no significant impact on photosynthetic characteristics of young fully expanded flag leaves (Table 1 and Figure 2). The maximum rates of photosynthesis on an area basis achieved at saturating CO_2_ partial pressure and high light measured at 25°C for high-temperature-grown plants were not different from plants grown at control temperatures. Nevertheless, an increase in the thermal optimum of photosynthesis was observed (Table 1). Total Rubisco activity and activation status were similar between treatments, however, heat-stressed plants displayed significant increases in PEPC activity and the PEPC:Rubisco ratio. The response of photosynthetic rate to CO_2_ concentration changes at high light, and to light level changes at ambient CO_2_ was indistinguishable from plants grown at control temperatures (Figure 2). Similarly, the photosynthetic carbon assimilation response at varying temperatures did not differ between heat-stressed and non-stressed plants (Supplementary Figure 2). Furthermore, bundle sheath leakiness (φ) determined from online stable isotope discrimination was unchanged across measurement temperatures, which indicated CCM function was not perturbed between treatments (Supplementary Figure 2).

**Figure 2:**
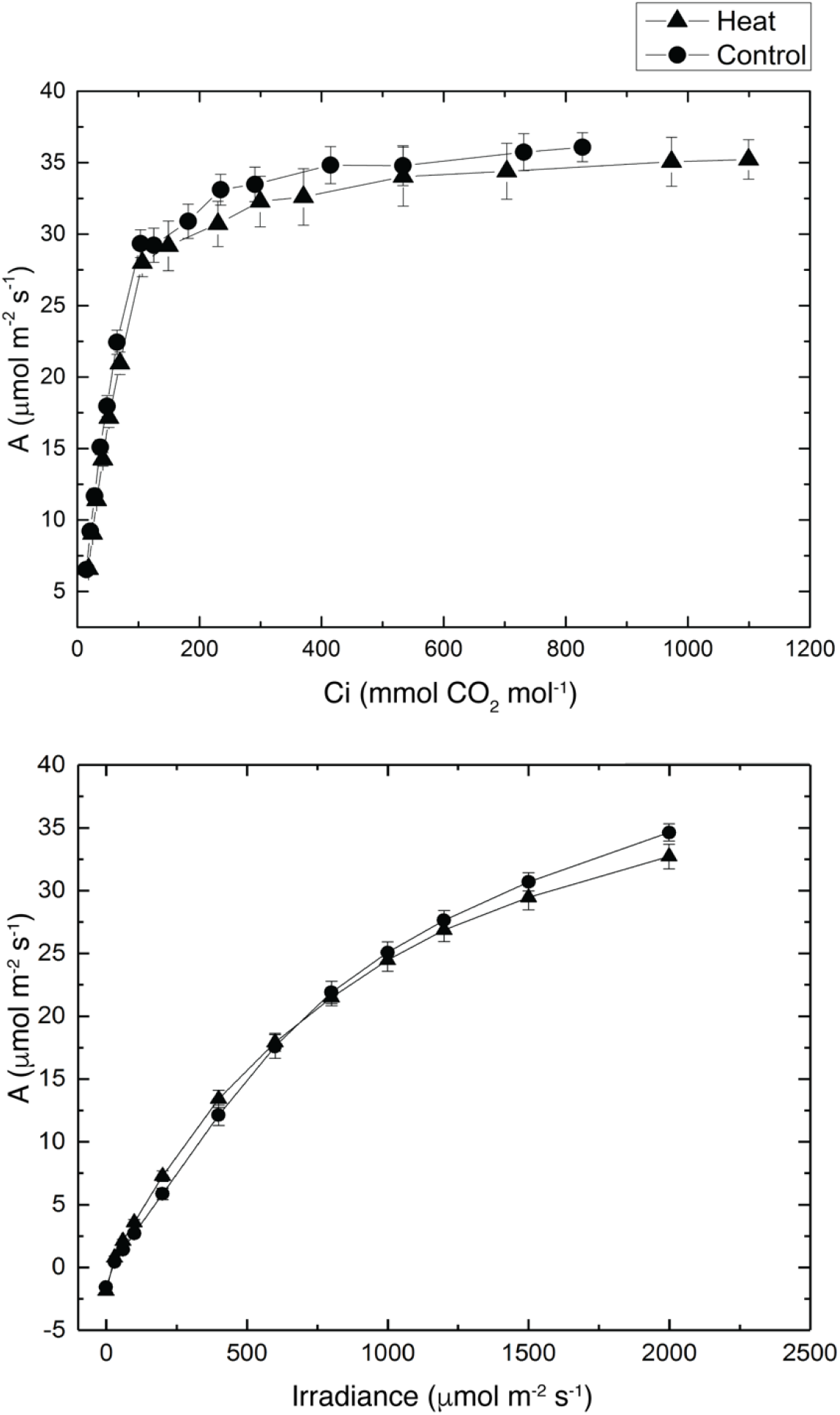
A: The CO_2_ assimilation rate (A) of the flag leaf in response to increasing concentrations of CO_2_, i.e. the A/Ci curve, of the control and heat-stressed plants. Measurements were taken at 25 °C and irradiance of 2000 μmol m^-2^s^-1^. The initial slopes of the A/Ci curves for each replicate plant were calculated separately by fitting a linear regression line through the first five data points. The initial slope for the control plants was 0.34 ± 0.01, and 0.31 ± 0.01 for the heat-stressed plants (p = 0.18). (N = 4). B: The response of CO_2_ assimilation rate to changing light intensity in the control and heat-stressed plants. Measurements were performed at ambient CO_2_ concentration of 380 μbar and 25 °C. Closed circles represent means of 4 biological replicates of plants grown at 28 °C day/ 22 °C night. Closed triangles represent means of 4 biological replicates of plants grown at 42 °C day/ 32 °C night.

**Table 1:** Summary of biochemical characteristics of the leaf of S. viridis plants grown at 28 °C day/22 °C night or 42 °C day/32 °C night. N=4 for each parameter.

|  | Growth Conditions |  | p-value<br>(t-test) |
| --- | --- | --- | --- |
|  | 28 °C / 22 °C | 42 °C / 32 °C |  |
| **A, at 25 °C ($\mu\text{mol m}^{-2} \text{s}^{-1}$)** | 28.81 ± 3.53 | 23.55 ± 1.62 | ns |
| **A, at growth temperature** | 32.19 ± 3.50 | 30.09 ± 2.12 | ns |
| **T<sub>opt</sub> for A (°C)** | 34.86 ± 0.70 | 37.25 ± 0.44 | 3.70E-02 |
| **Rubisco activity (initial)**<br>**($\mu\text{mol CO}_2 \text{m}^{-2} \text{s}^{-1}$)** | 16.23 ± 2.32 | 20.29 ± 3.07 | ns |
| **Rubisco activity (activated)**<br>**($\mu\text{mol CO}_2 \text{m}^{-2} \text{s}^{-1}$)** | 22.61 ± 1.74 | 27.82 ± 1.00 | ns |
| **Rubisco activation %** | 71.86 ± 8.26 | 72.82 ± 10.55 | ns |
| **PEPC activity ($\mu\text{mol CO}_2 \text{m}^{-2} \text{s}^{-1}$)** | 187.82 ± 6.96 | 301.44 ± 10.11 | 7.60E-04 |
| **PEP:Rubisco** | 8.36 ± 0.36 | 10.84 ± 0.05 | 2.50E-03 |
| **Chlorophyll a+b ($\text{mmol m}^{-2}$)** | 0.70 ± 0.03 | 0.68 ± 0.06 | ns |
| **Chlorophyll a/b** | 6.56 ± 0.26 | 5.29 ± 0.35 | 2.00E-02 |
| **Nitrogen ($\text{mmol m}^{-2}$)** | 82.49 ± 6.17 | 116.84 ± 6.99 | 1.10E-03 |
| **C:N** | 21.17 ± 4.29 | 12.21 ± 0.64 | 8.10E-03 |

### Transcript, protein and metabolite profiles

#### The C_4_ cycle

To further investigate the effect of long-term heat stress on the functioning of the C_4_ cycle, the expression of genes involved in the C_4_ pathway was examined using both RNASeq and quantitative proteomics methods, coupled to metabolite measurements by GC-MS. Levels of mRNA transcript (T), protein (P) and metabolites in leaves of the control and heat-stressed plants are shown in the color-coded diagram in Figure 3. Numbers in the text box next to enzyme abbreviation represent the ratio between heat-stressed vs control plants in log2 scale. Although transcript levels of core C_4_ cycle enzymes (carbonic anhydrase, CA; phosphoenolpyruvate carboxylase, PEPC; NADP-malate dehydrogenase, NADP-MDH; Rubisco large subunit, RBCL, Rubisco small subunit, RBCS; and pyruvate orthophosphate di-kinase, PPDK) were significantly lower under heat stress, in most cases there were no significant reductions in the amount of proteins, except for NADP-ME and RBCS (Figure 3). Similarly, the transcript levels of transporters for C_4_ pathway metabolites (OMT1, DIT1, DCT2/DIT2, MEP3a and b, PPT, TPT) (Weber and von Caemmerer, 2010; Schlüter et al., 2016) were significantly lower. A significant reduction at the protein level was only observed for the dicarboxylate transporter DCT/DIT2, which transports malate into the BS chloroplast for decarboxylation (Weissmann et al., 2016). There was a significant buildup of malate in the leaves of heat-stressed plants although it is unclear in which cell type or subcellular compartment the malate accumulated.

**Figure 3.**
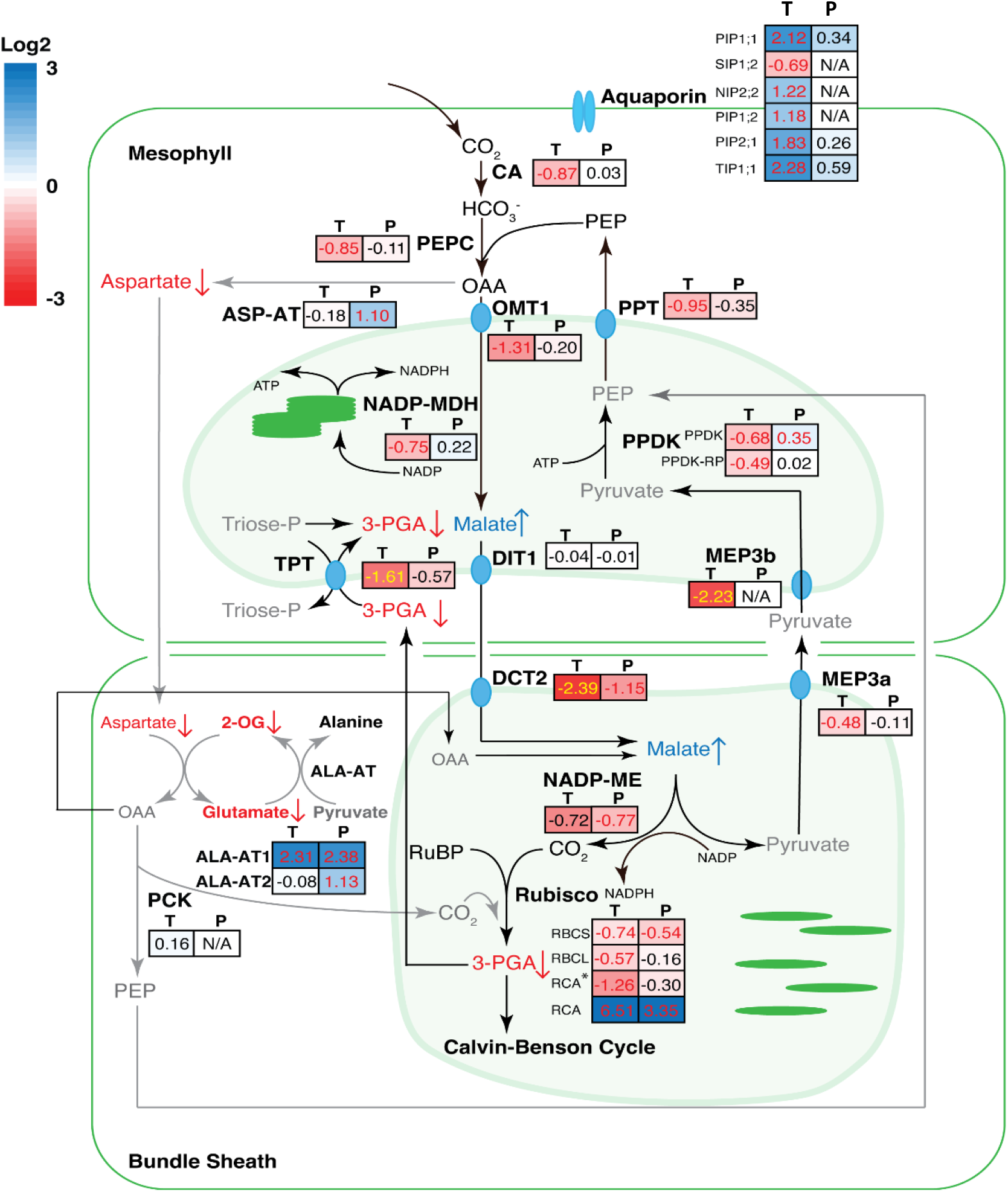
The response in expression of genes involved in the C_4_ pathway at the transcript and protein level, and the associated changes in metabolites to heat stress treatment. For each gene, the transcript and protein fold-changes (Heat Vs Control) in log2 scale are shown in coloured boxes, where “T” and “P” designate transcript and protein levels, respectively. The colour scale for transcript and protein log_2_ fold-changes can be found on the top left corner. Numbers in red (or yellow if background is dark red) indicate the fold-changes are significant with p < 0.05. Metabolites detected by GC-MS are shown in red (with down arrow), blue (with up arrow) or black, indicating significant (p<0.05) decrease, significant increase or no significant change, respectively. Metabolites not detected were shown in grey. Relative metabolite levels are shown in Supplementary Data File 2. Abbreviations used: CA, carbonic anhydrase; PEPC, phosphoenolpyruvate carboxylase; ASP-AT, aspartate amino transferase; OMT1, oxoglutarate:malate antiporter; NADP-MDH, NADP-dependent malate dehydrogenase; PPDK, pyruvate orthophosphate dikinase; DIT1, dicarboxylate transporter 1; TPT, triose-phosphate translocator; NADP-ME, NADP-dependent malic enzyme; Rubisco, ribulose bisphosphate carboxylase/oxygenase; ALA-AT, alanine aminotransferase; PCK, phosphoenolpyruvate carboxykinase.

Some NADP-ME type C_4_ plants such as *Zea mays* and *Flaveria bidentis* produce a significant amount of aspartate in the mesophyll cells as the C_4_ metabolites for translocation of carbon to the BS cells, where it can be converted to oxaloacetate and decarboxylated by PEP carboxykinase (PEPCK), or reduced to malate and decarboxylated by NADP-ME (Furbank, 2011). In our experiment, the expression of aspartate aminotransferase (ASP-AT), which converts OAA into aspartate, was significantly upregulated at the protein level under heat stress but PEPCK transcript was undetectable (Figure 3; Supplementary Data File 1 - Table 1). Concordantly, the expression of two alanine aminotransferases (ALA-AT) were also upregulated significantly at the protein level. The levels of aspartate, 2-OG, and glutamate significantly reduced in the heat-stressed plants (Figure 3). The heightened flux in the reaction pathway to producing PEP from aspartate suggests that under heat stress, aspartate is an important C_4_ acid of the C_4_ cycle, potentially allowing the products of PEP carboxylation to move to the bundle sheath cells under conditions where the mesophyll chloroplast stroma is becoming more oxidized and less able to reduce OAA to malate. Interestingly, we also observed a concurrent highly significant decrease in the level of beta-alanine (Supplementary Data File 2), a metabolite long suspected (but so far not proven) to be derivable from aspartate in plants that has previously been observed to *increase* under heat stress in divergent species (Parthasarathy et al., 2019) and linked to differential tolerance against abiotic stresses, including heat stress, in plants (Cross et al., 2004; Fouad and Altpeter, 2009; Glaubitz et al., 2017; Devnarain et al., 2019; Parthasarathy et al., 2019).

#### The Calvin-Benson-Bassham cycle

The effect of heat stress on the expression of genes involved in the Cavlin-Benson-Bassham (CBB) and the photorespiratory cycles was also examined. The complexity of the traffic of 3-C sugar phosphates between the mesophyll and bundle sheath cells of C_4_ plants (von Caemmerer and Furbank, 2016) and the absence of cell-specific data in this study require some assumptions in interpreting data. The pathway diagram shown in Figure 4 is based on cell-specific gene expression data in the two cell types (John et al., 2014) where each gene is placed in the location where it is predominantly expressed. All of the genes involved in the CBB cycle showed significant reduction at the transcript level in the heat-stressed plants compared to the control (Figure 4). However, at the protein level, only phosphoribulokinase (PRK) reduced significantly at high temperature, while other enzymes were unchanged.

**Figure 4.**
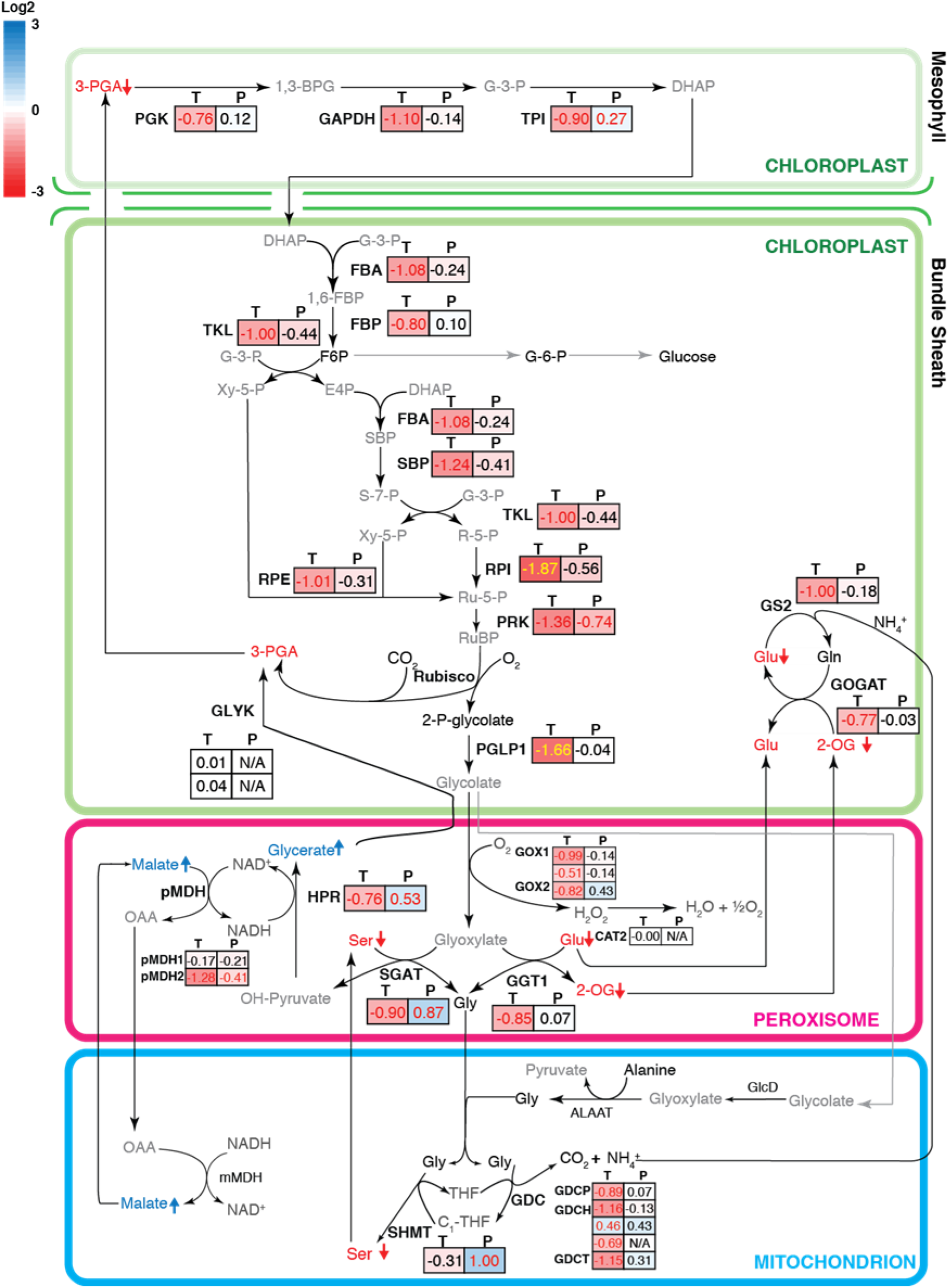
The response in expression of genes involved in the Benson-Calvin cycle and the photorespiratory cycle at both the transcript and protein level, and the associated changes in metabolites involved in the pathways to heat stress. The colour codes of the diagram are the same as described in Figure 3. Abbreviations used: PGK, phosphoglycerate kinase; GAPDH, glyceraldehyde-3-phosphate dehydrogenase; TPI, triose phosphate isomerase; FBA, fructose-bisphosphate aldolase; FBP, fructose bisphosphatase; TKL, transketolase; SBP, sedoheptulose-1,7-bisphosphatase; RPE, ribulose-phosphate3 epimerase; RPI, ribose-5-phosphate isomerase; PRK, phosphoribulose kinase; PGLP1, phosphoglycolate phosphatase1; GOX, glycolate oxidase; SGAT, serine:glyoxylate aminotransferase; GGT, glutamate:glyoxylate aminotransferase; SHMT, serine hydroxymethyltransferase; GDC, glycine decarboxylase; mMDH, mitochondrial malate dehydrogenase; pMDH, peroxisomal malate dehydrogenase; HPR, hydroxypyruvate reductase; CAT, catalase; GS, glutamine synthetase; GOGAT, ferrodoxin-dependent glutamate synthase; GLYK, glycerate kinase.

Notably, both transcript and protein levels of an isoform of RCA with an C-terminal extension (Supplementary Figure 3) was strongly induced and almost exclusively expressed under heat stress (the transcript increased ∼90-fold and the protein increased ∼10-fold under heat treatment). It is known that activase is a key heat sensitive step in photosynthesis (Law and Crafts-Brandner, 1999; Salvucci and Crafts-Brandner, 2004). The C-terminal tail has been suggested to play a role in Rubisco activation under heat stress in both wheat and maize, as a longer RCA isoform has been reported to be more heat stable (Crafts-Brandner and Salvucci, 2002; Ristic et al., 2009).

#### The photorespiratory cycle

Although C_4_ plants have a largely diminished level of photorespiration, the genes involved in the photorespiratory cycle were still highly expressed in *S. viridis*, based on the mean RPKM of the photorespiratory genes relative to genes involved in the Benson-Calvin cycle (Supplementary Data File 1 - Table 1). Although at the transcript level, expression of PGLP1, GOX, GGT1, SGAT, HPR, GDC subunits, pMDH2, Fd-GOGAT, and GS2 were reduced, the protein levels of SGAT, SHMT, and HPR significantly increased under heat stress (Figure 4). Accompanying these changes was a significant reduction in the level of serine, and a significant increase in the accumulation of glycerate (Figure 4; Supplementary Figure 5). This glycerate, produced in the photorespiratory cycle, needs to be converted back to 3-PGA to complete the cycle. It is uncertain whether changes in the level of GLYK would have any effect on the accumulation of glycerate as GLYK protein was not detected. Metabolites involved in the NH_4_^+^ assimilation part of the photorespiratory cycle, including glutamate and 2-oxoglutarate, also showed significant reduction in the heat-stressed plants (Figure 4).

### Expression of genes involved in chloroplast electron transport

Chloroplast electron transport consists of both linear and cyclic electron transport. The M and BS cells host linear and cyclic electron transport chain components, although cyclic electron transport occurs primarily in the BS cells in NADP-ME type C_4_ plants (Figure 5). The expression of genes in each protein complex in the electron transport chain was examined at the transcript and protein level. Transcripts encoding several components of the light harvesting complex (LHC) II and Photosystem II were expressed at a lower level under heat stress. In addition, the protein levels for psbO, psbP and psbQ, which are components of the oxygen evolving complex, also reduced significantly at the protein level (Figure 5). All of the subunits of Cytochrome b6f complex exhibited significant reduction in mRNA levels, with PETA and PETC also reduced significantly at the protein levels (Figure 5). Similarly, many genes encoding Light Harvesting Complex I and Photosystem I subunits showed significant decreases in expression at the transcript level, but changes in the corresponding proteins were mostly not significant. Subunits of ATP synthase all exhibited a similar magnitude of reduction by heat stress (except ATPD) at the transcript level, while protein abundance of ATPB, ATPC, ATPF, and ATPX also decreased significantly. The NDH complex, involved in cyclic electron transport activity (Johnson, 2011), showed a pronounced reduction of all subunits at the transcript and many of the subunits detected in the proteomics study were also significantly reduced. Expression of another protein involved in PSI cyclic electron transport, the PGR5 protein (Johnson, 2011), was also reduced at both the transcript and protein level under high temperature.

**Figure 5.**
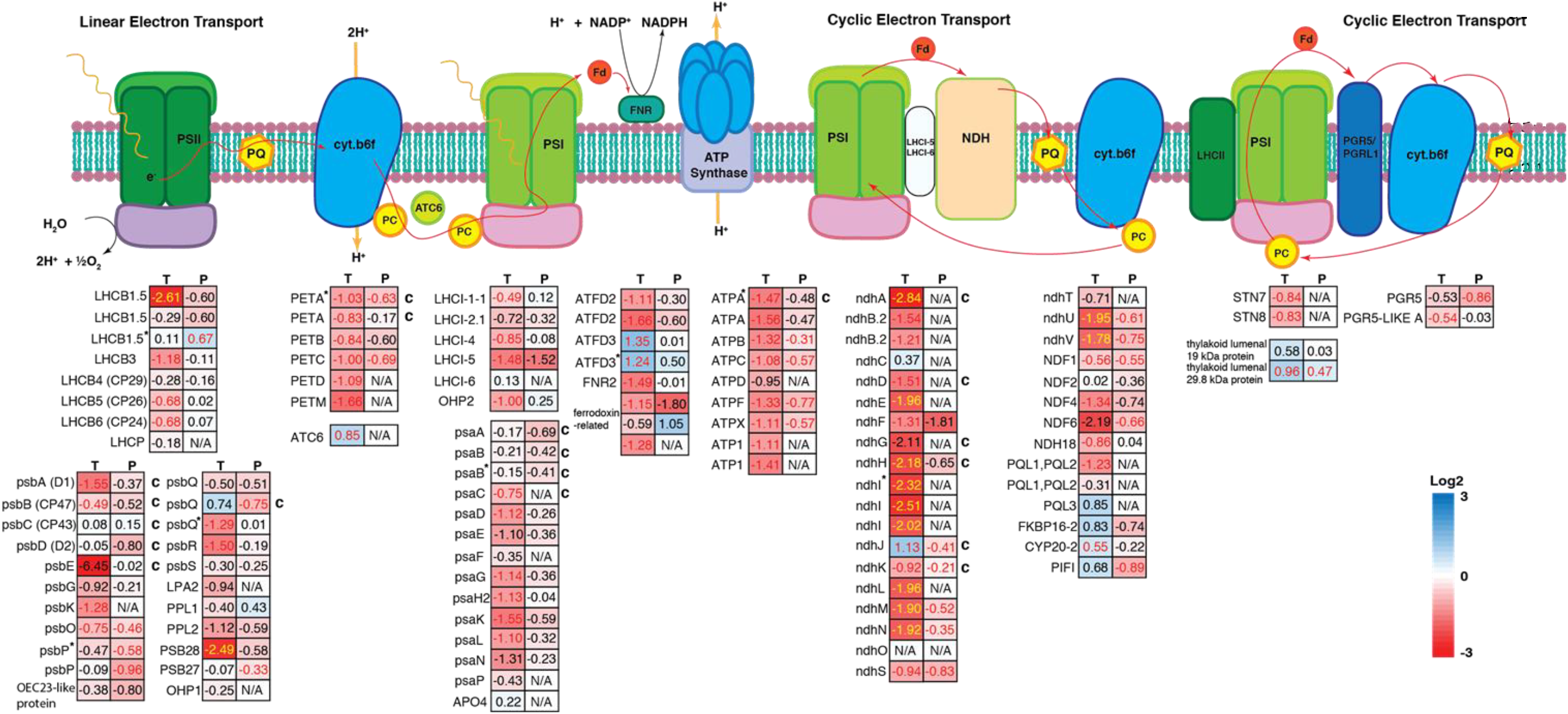
Expression of chloroplast electron transport genes under heat stress treatment. Schematics of protein complexes of the chloroplast electron transport chain located on the thylakoid membrane are drawn. Linear electron transport includes the Photosystem II, Cytochrome b6f, Photosystem I, ATP synthase complexes. There are two possible modes of cyclic electron transport which involve Photosystem I, NDH complex or PGR5 protein. The expression level of the subunits of each protein complex is shown below the schematic diagram. Numbers shown are the log2 transformed fold changes between the heat-stressed plants and control plants. The left column shows the expression level of transcripts and the right column shows protein level. When multiple isoforms of the same subunit exist, an asterisks mark * is used to designate the most highly expressed isoform. A letter “c” is written next to genes encoded by the chloroplast genome. The color scale can be found on the bottom right corner.

Of note is the concerted reduction of expression of different subunits of the same complex. For example, expression of the subunits of Cytochrome b6f, ATP synthase, and NDH complex all showed similar levels of reduction at the transcript level. This suggests that the expression of these subunits of the same complex might be under the regulation of the same transcription regulon. Despite the global reduction in expression in the components of chloroplast electron transport chain, electron transport capacity was maintained in the heat-stressed plants (Figure 2B).

### Soluble sugars and starch levels

The leaf starch and sugar contents in 4-week-old *S. viridis* plants grown at 28 °C (control) or 42 °C (heat-stressed) were measured in leaf samples collected at mid-day. On a leaf area basis, heat-stressed plants accumulated significantly smaller amounts of starch compared to the control counterparts (Figure 6A). This result was further confirmed by TEM images of the leaves, showing reduced accumulation of starch granules inside chloroplasts in the heat-stressed plants compared to the control (Supplementary Figure 4).

**Fig 6:**
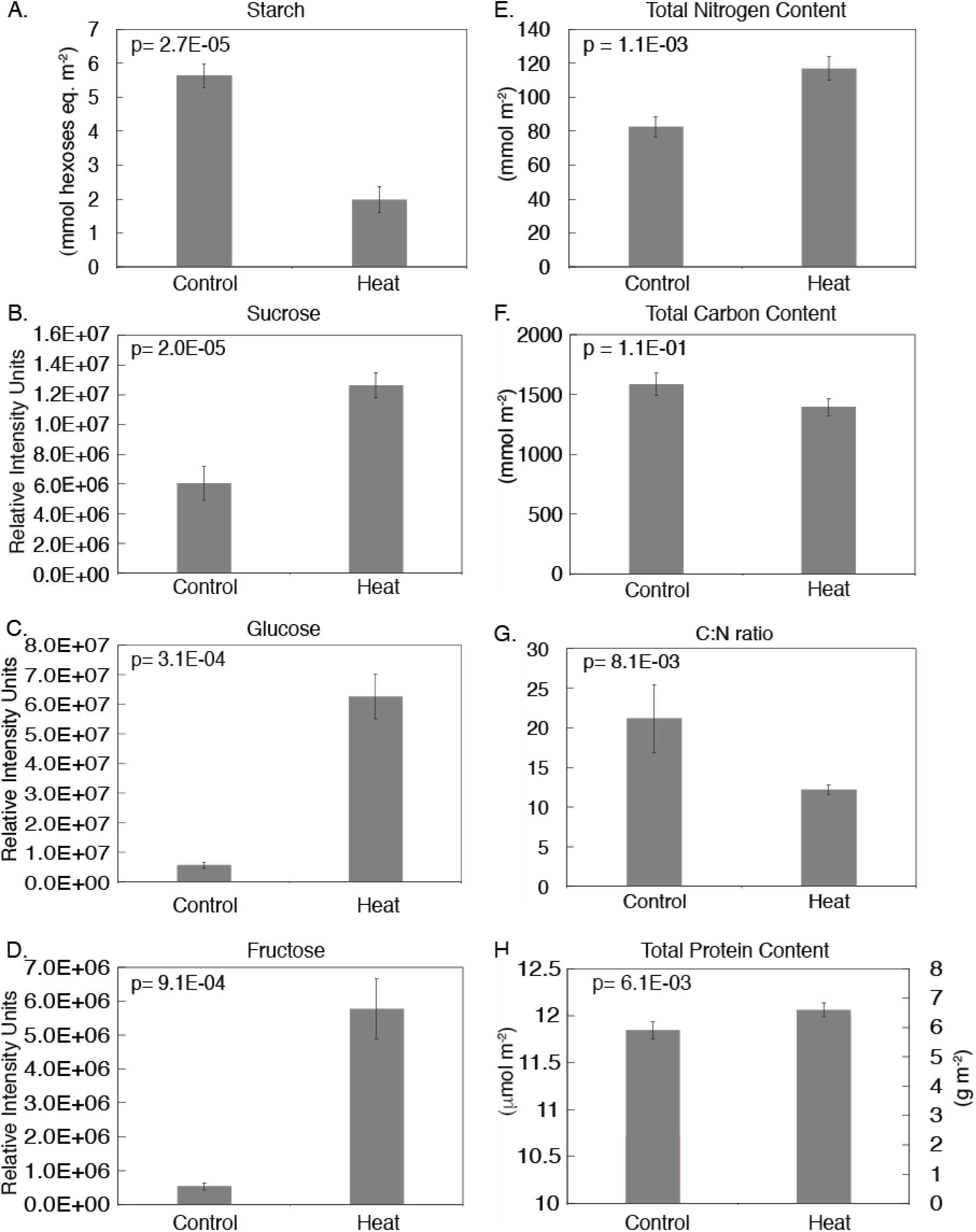
The levels of starch, soluble sugars (sucrose, glucose and fructose), total nitrogen, protein and chlorophyll content, and carbon: nitrogen (C: N) ratio in the control and heat-stressed plants are shown in bar graphs, with the standard error of mean (SEM) obtained from measurements of 5-6 biological replicates. The levels of soluble sugars are shown as relative concentrations, in units of relative intensity, which represents the signal intensity of the derivatised compounds on a GC-MS.

The relative levels of leaf soluble sugars were measured by GC-MS metabolomics, and are shown in units of relative intensity in Figure 6B-D. The fold-changes of all the metabolites measured by GC-MS and their p-values can be found in Supplementary Figure 5. The heat-stressed plants accumulated significantly higher levels of sucrose, glucose and fructose in leaves (Figure 6B-D). The level of glucose-6-phosphate did not change significantly in the heat-stressed plants compared to the controls and there was only a small but significant increase in the level of fructose-6-phosphate (Supplementary Figure 5). The levels of trehalose, raffinose and proline increased strongly in the heat-stressed plants (Figure 7 A-C). In addition, a number of other soluble sugars (including cellobiose, gentiobiose, maltose, arabinose, ribose, xylulose and mannose) and sugar alcohols (including D-threitol, erythritol, sorbitol and xylitol) also showed significantly increased levels in the heat-stressed plants (Supplementary Figure 5).

**Figure 7:**
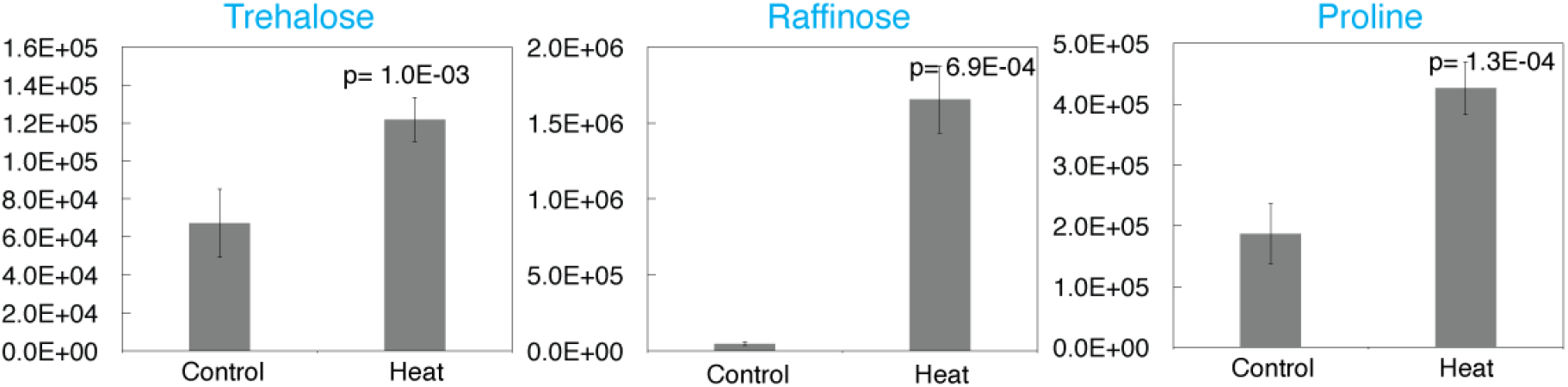
The relative levels of trehalose, raffinose and proline in the control and heat-stressed plants, as determined by GC-MS metabolomics, are shown in bar graphs. Values represent the integrated peak area of the m/z channel used to quantify each analyte after normalization to the peak area of the ribitol internal standard in the same sample. Peak area of the quantifier ion for each metabolite, in arbitrary signal intensity unit. These values are only comparable between measurements of the same metabolites, not between different metabolites. Compounds significantly increased (p<0.05) are highlighted in blue, significantly decreased are highlighted in red. Student t-test were performed. N=5.

Interestingly, while leaf protein levels increased only slightly in heat stressed leaves, nitrogen content increased and the carbon to nitrogen ratio of the tissue under heat stress declined by almost 50% (Figure 6 G,H).

Taken together, these results showed the significant accumulation of a wide range of soluble sugars and their derivatives in the heat-stressed *S. viridis* plants, whereas the accumulation of transitory starch significantly reduced under heat.

### Expression of genes encoding enzymes of carbohydrate metabolism

Given the large reprogramming of sugar and starch metabolism during heat stress shown in Figures 6 and 7, expression of genes encoding enzymes in starch and sugar biosynthesis were examined (Figure 8). For simplicity of the diagram, Figure 8 used the proposed cell localization of carbohydrate metabolism for maize (Lunn and Furbank, 1999), with starch synthesis only shown inside the BS cell and sucrose synthesis/breakdown only shown inside the M cell.

**Figure 8:**
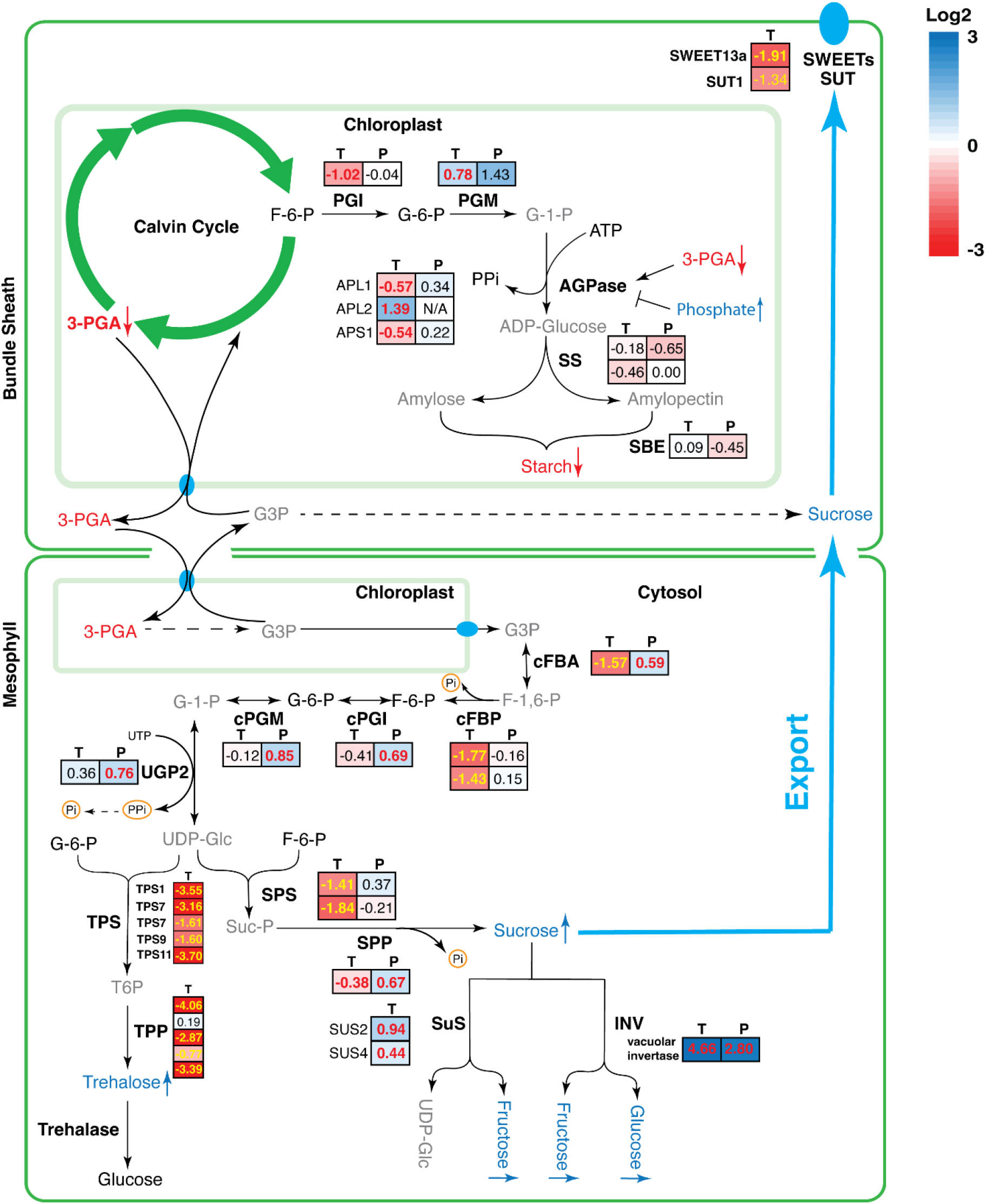
Changes in the expression of genes encoding for proteins involved in starch synthesis, sucrose synthesis/breakdown and trehalose metabolism. For each gene, the transcript and protein fold-changes (Heat Vs Control) in log2 scale are shown in coloured boxes, where “T” designate the change in transcript level and “P” designate protein level. The colour scale can be found on the top right corner. Numbers in red (or yellow if background is dark red) indicate the fold-changes are significant with p < 0.05. Metabolites with levels that increased or decreased markedly (with p<0.07) are shown in blue or red, respectively, while metabolites that were detected but did not change significantly are shown in black. Metabolites that were not measured are shown in grey.

The genes involved in starch synthesis, AGPase large subunit (APL1) and small subunits (APS1), SS (Starch Synthase) and SBE (Starch Branching Enzyme) did not show significant changes at the level of protein abundance in the heat-stressed plants, although the transcript levels of APL1 and APS1 significantly decreased (Figure 8).

The genes involved in sucrose synthesis, including cFBA (cytosolic Fructose Bisphosphate Aldolase), cPGI (cytosolic Glucose-6-phosphate Isomerase), cPGM (cytosolic Phosphoglucomutase), UGP2 (UDP Glucose Pyrophospharylase) and SPP (Sucrose Phosphate Phosphatase) all increased significantly in terms of protein abundance. However, the transcript abundance of cFBA, cFBP and SPS (Sucrose Phosphate Synthase) significantly decreased under heat (Figure 8). The expression of SUS (Sucrose synthase) and vacuolar INV (Invertase) significantly increased under heat, and the protein abundance for vacuolar INV also showed a significant increase. Metabolomics data (Figure 7) showed a significant increase in sucrose, glucose and fructose levels in the heat-stressed plants consistent with up-regulation of sucrose synthesis and its breakdown into hexoses under heat. The hexose sugars might be accumulating inside the vacuole as vacuolar INV was significantly up-regulated.

The synthesis of trehalose-6-phosphate (T6P) has been shown to be closely correlated with the amount of sucrose in the cells (Paul, 2007; Figueroa and Lunn, 2016). In the current experiment, we could not measure the level of T6P, but the level of trehalose significantly increased in the heat-stressed plants. However, transcript levels for various TPS (Trehalose-6-phosphate Synthase) and TPP (Trehalose-6-phosphate Phosphatase) isoforms have all decreased significantly and the protein products were not detected in this experiment (Figure 8).

In concert with the significant accumulation of raffinose (∼35-fold increase compared to control) and galactinol (∼4-fold increase) in response to high temperature, the expression of galactinol synthase 1 (GolS1), raffinose synthase and stachyose synthase were significantly up-regulated at the transcript level (Supplementary Figure 6). The expression of GolS1 & 2 and stachyose synthase are preferentially localized to M cells while one raffinose synthase isoform preferentially localizes to BS cells and the other two isoforms do not show cell type preference in the published cell specific expression data sets of John et al. (2014).

### Changes in hormones under heat

The visible phenotype of dwarfing / stunting of all plant parts in *Setaria* grown under high temperature is striking (Figure 1). To examine this phenotype in more detail, leaf hormone levels were profiled in heat stressed and control plants (Figure 9). The levels of SA (salicylic acid), JA (jasmonic acid) and IAA (indole-3-acetic acid) did not change significantly, nor did the level of phenylacetic acid (PAA), an auxin analogue in plants (Figure 9). However, the level of ABA increased markedly to about 20-fold over the control levels in the heat-stressed plant. Although the level of IAA did not change, there was massive accumulation of IAA-Aspartate amino conjugate in the heat-stressed plants while IAA-Aspartate amino conjugate is not detectable in the control plants. This compound is proposed to be an inactive reservoir for IAA (Ruiz Rosquete et al., 2012) or in one study a potent signal during abiotic stress (Ostrowski et al., 2016). In addition, there was also a significant accumulation of IAA-Alanine amino conjugate in the heat-stressed plants.

**Figure 9:**
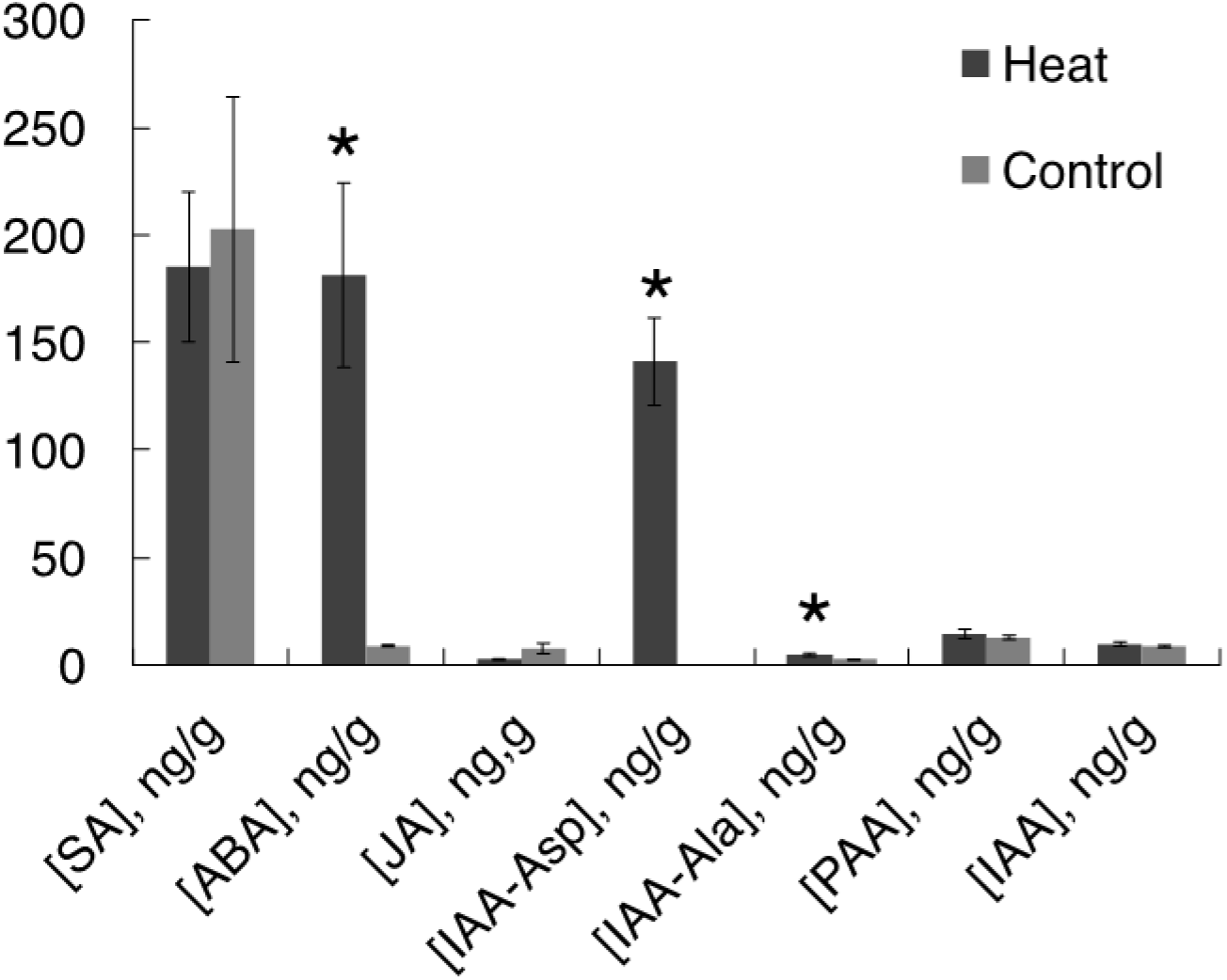
Measurements of hormone levels using LC-MS. SA, salicylic acid; ABA, abscisic acid; JA, jasmonic acid; IAA, indole-3-acetic acid (i.e. auxin); IAA-Asp, auxin-aspartate amino conjugate; IAA-Ala, auxin-alanine amino conjugate; PAA, phenylacetic acid. Asterisks indicate a significant difference at p < 0.05. Number of samples measured for each hormone and for each treatment is 4 or 5.

**Figure 10:**
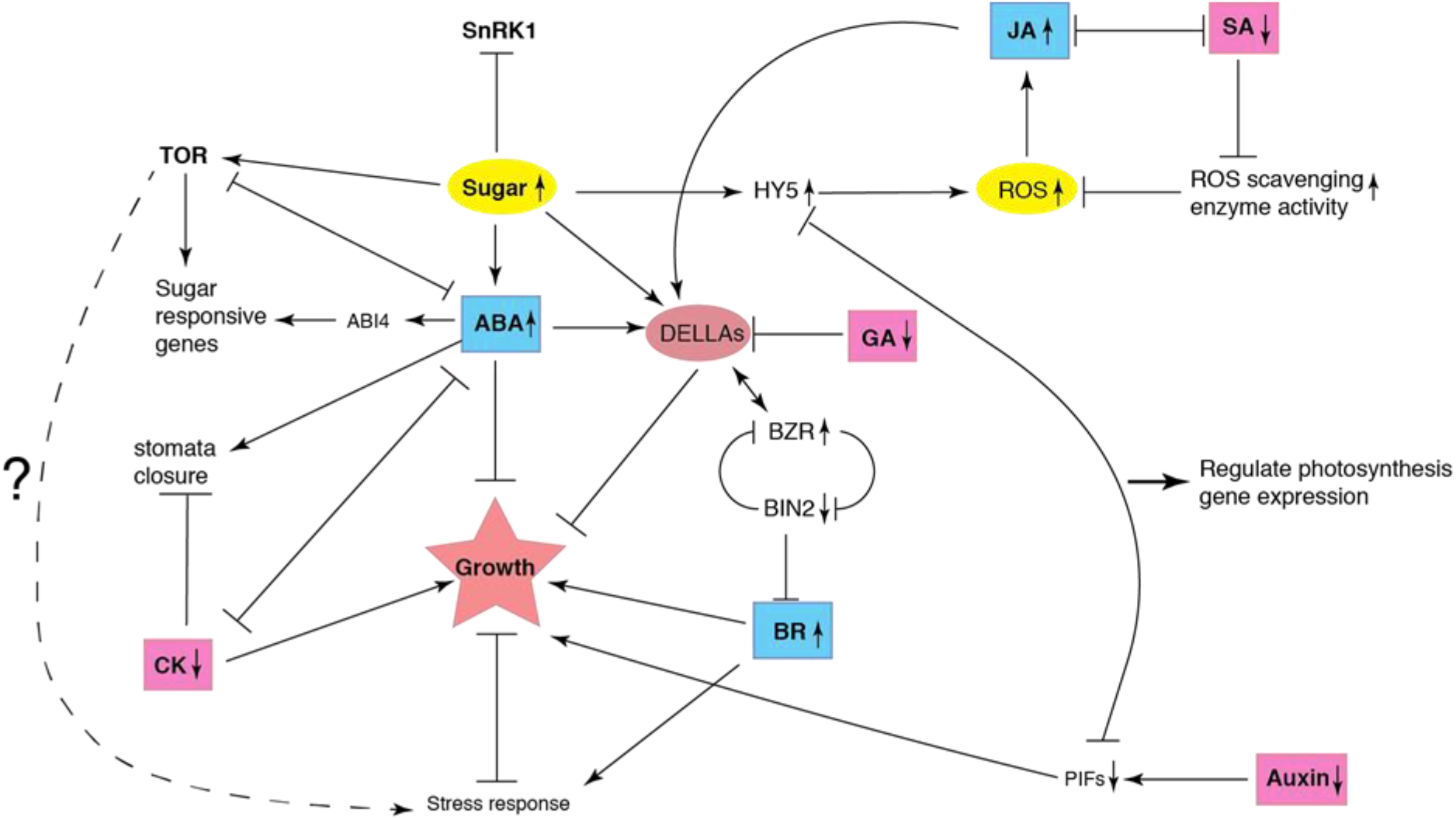
A hypothetical model of the regulation of growth and heat stress response by phytohormones and sugar signaling in S.viridis. Arrows indicate a positive relationship where one component may stimulate/stabilize the other component. Lines with a blunt end indicate an inhibitory relationship, where one component negatively interacts /regulates the other component. Abbreviations used: ABA, abscisic acid; CK, cytokinin; GA, gibberellin; BR, brassinosteroid; SA, salicylic acid; JA, jasmonic acid; ROS, reactive oxygen species; TOR, target of rapamycin; SnRK1, sucrose non-fermenting related kinase 1; HY5, long hypocotyl 5; BZR, brassinosteroid positive regulator related; BIN2, brassinosteroid insensitive 2; ABI4, ABA insensitive 4; PIF, phytochrome interacting factor.

## Discussion

### Effects of heat stress on photosynthesis and growth

The lack of effect on the rate of photosynthesis in response to CO_2_ and irradiance and on levels of photosynthetic proteins in the leaves of plants grown for 2 weeks at 42/32°C was unexpected (Figures 2 and 3). The marked dwarfing observed (Figure 1) suggests that carbon supply might be severely limiting. The thermal response of photosynthesis shown in Supplementary Figure 2 suggests that C_4_ photosynthesis in *S. viridis* was relatively insensitive to the exposure of elevated temperatures, unlike other reports for C_4_ species. For example, CO_2_ assimilation rates dropped sharply at temperatures above 38 °C in *Zea mays* (Crafts-Brandner and Salvucci, 2002). In sorghum, photosynthesis declined after 3 hours’ incubation at 40 °C (Yan et al., 2011).Similarly for *S. viridis* plants exposed to 40°C for 4 hours, reductions in CO_2_ assimilation were observed (Anderson et al., 2021). Several C_4_ species (monocot *Panicum coloratum* and *Cenchrus ciliaris, and* dicot *Flaveria bidentis*) grown at moderately high temperatures (35 / 30 °C) also exhibited reduced photosynthetic capacity compared to plants grown at control temperatures (25 / 20 °C) (Dwyer et al., 2007). Other crop species such as cotton and wheat also exhibited significant reduction of photosynthesis at temperatures beyond 40 °C (Law and Crafts-Brandner, 1999). However, in the present study, both *S. viridis* plants grown at control (28 / 22 °C) or high temperatures (42 / 32 °C) suffered relatively little loss of photosynthetic activity at 45 °C (Supplementary Figure 2).

Major points of high temperature sensitivity have been identified in the photosynthesis pathway, including chloroplast electron transport components affected under high temperature and high light (Nash et al., 1985; Takahashi et al., 2004; Sharkey, 2005; Allakhverdiev et al., 2008a), and inhibition of caused by RCA protein turnover, denaturation and loss of activity (Sharkey et al., 2001; Salvucci and Crafts-Brandner, 2004; Salvucci and Crafts-Brandner, 2004). However, despite transcriptome analyses identifying levels of many photosynthetic transcripts were reduced at high temperatures, this was only reflected in the proteomics profile in a small number of cases (Figures 3-5). Lack of correlation between transcript levels and protein levels or non-linear relationships between the two are not uncommon in the plant literature (Hajduch et al., 2010; Ponnala et al., 2014; Vélez-Bermúdez and Schmidt, 2014). Some possible explanations for this lack of correlation are discussed below. First, mature, fully expanded leaf blades were used for this study. Depending on the stability of individual photosynthetic proteins (particularly under heat stress) and their developmental timing of synthesis, peak transcriptional activity may occur very early in leaf development and not coincide with protein level in mature tissue (Ponnala et al., 2014). Second, control of protein levels of some photosynthetic proteins, in particular those forming complexes in the chloroplast thylakoid, are believed to be controlled post-transcriptionally (Choquet and Wollman, 2002; Eberhard et al., 2002; Park et al., 2012; Floris et al., 2013; Fankhauser and Aubry, 2016; Zoschke and Bock, 2018; Perdomo et al., 2021), and are often extremely long-lived proteins (Adam, 1996; Bienvenut et al., 2011; Nelson and Millar, 2015).

The lack of obvious physiological effects on photosynthesis after 42 °C growth in these experiments suggests that the photosynthetic machinery of *S. viridis* had become thermally acclimated to 42 °C. Evidence for heat adaptation at 42 °C is supported by an increased T_opt_ for photosynthesis (Table 1), maintenance of photosynthesis rates at the high growth temperature and higher photosynthesis rates at 45 °C in the acclimated plants (Supplementary Figure 2), maintenance of photosynthesis rates at the new T_opt_ and maintenance of photosynthetic capacity measured at a basal temperature of 25 °C (Figure 2 and Supplementary Figure 2). These changes fulfilled all the criteria outlined by Way and Yamori (2014) of thermal acclimation of photosynthesis and suggests that a positive adjustment has occurred to optimize the plant response to heat stress. In this regard, there appears to have been some adaptive responses for a subset of photosynthetic proteins and a response to heat treatment in photosynthetic metabolite profiles (Figures 3 and 4).

Notably, expression of chloroplast membrane transporters involved in C_4_ photosynthesis was suppressed at high temperatures at the transcript level and in the case of DCT2/DIT2 also at the protein level (Figure 3). This may have been the cause for an increase in foliar malate levels at high temperature as DCT2/DIT2 is believed to move malate into the bundle sheath chloroplast (Weber and von Caemmerer, 2010). However, photosynthetic malate probably only makes up a relatively small proportion of total leaf malate pools (Hatch, 1979; Meister et al., 1996). In concert with a rise in malate levels, aspartate levels fell and levels of aspartate and alanine aminotransferase enzymes increased under heat (Figure 3). This could be indicative of a shift toward aspartate formation from OAA in the mesophyll and its utilization in the bundle sheath to regenerate malate for decarboxylation as seen in the NADP-ME type C_4_ maize and Flaveria (Meister et al., 1996; Furbank, 2011; Arrivault et al., 2016).

A shift toward aspartate as the translocated C_4_ metabolite under heat would likely be triggered by the strong induction of pathways implicated in combating oxidative stress (Supplementary Figure 7). The reduction of oxidized glutathione and dehydroascorbate during detoxification of H_2_O_2_ occurs exclusively in the mesophyll compartment (Doulis et al., 1997) which would result in strong competition for reducing power in the mesophyll chloroplasts and reduced capacity for OAA reduction (Turkan et al., 2018). Furthermore, it can be hypothesized that there is a build-up of foliar nitrate under heat stress would correspond to the observed increase in leaf nitrogen and down-regulation of the gene expression in the nitrate assimilation pathway and contribute to the lack of reducing power in the chloroplast (Supplementary Figure 8). Nitrate and nitrite reductase are localized to the mesophyll cells in NADPME-type C_4_ grasses (Moore and Black Jr, 1979) consistent with competition for reducing power between OAA reduction, oxidative stress protection and nitrogen metabolism in this compartment during heat treatment.

While adaptive responses in the C_3_ photosynthetic machinery and photorespiratory pathways of *Setaria* under high temperature were not generally large, the response of RCA expression (Figure 3) was marked. RCA has been identified as a key point of heat sensitivity in the photosynthetic machinery of both C_3_ and C_4_ plants (Law and Crafts-Brandner, 1999; Crafts-Brandner and Salvucci, 2002; Salvucci and Crafts-Brandner, 2004; Salvucci and Crafts-Brandner, 2004; Busch and Sage, 2017; Scafaro et al., 2019; Degen et al., 2021). It has been proposed that the C-terminal extension is responsible for heat stability of Rubisco activase and the induction of the longer Rubisco activase isoform in heat-stressed *Setaria* supports this hypothesis (see Figure 3 and Supplementary Figure 3; Anderson et al. (2021)).

### Carbon partitioning and export

One of the largest metabolic effects of high temperature growth of *Setaria* was on foliar carbohydrate levels (Figures 6 and 7). A large shift occurred away from starch and in favour of high soluble sugar levels in leaves harvested mid-way through the photoperiod. Starch levels declined by almost two thirds under heat treatment, sucrose more than doubled and glucose and fructose levels increased more than 10-fold (Figure 6). This was in part due to changes in levels of key enzymes in carbohydrate synthesis but presumably also due to reduced sucrose export, as SWEET13 expression was severely reduced under heat stress (Figure 8). While compartmentation of these carbohydrates between M and BSC was not determined, nor was sub-cellular location, these are massive changes in carbon allocation with likely far reaching impact on osmotic regulation and sugar signaling pathways (Li and Sheen, 2016; Saddhe et al., 2021). Examination of the transcript and proteome profiles performed here shed some light on the mechanisms underlying these changes in carbohydrate levels.

The control on the rate of sucrose synthesis is thought to be mainly exerted through regulations on the activities of cFBPase and SPS in both C_3_ and C_4_ plants (Lunn and Furbank, 1999). Here the levels of cFBA, cPGI, cPGM, UGP2 and SPP proteins, which are all part of the sucrose synthesis pathway, significantly increased (Figure 5), which is consistent with an increase in sucrose biosynthesis. However, there was no change in the amount of cFBPase and SPS proteins and their transcript abundances significantly decreased (Figure 8). This may indicate the importance of post-translational regulation of cFBPase and SPS activity.

The increased accumulation of sucrose in the leaves of heat-stressed plants could additionally be caused by reduced phloem loading. Phloem loading of sucrose is thought to occur by export of sucrose into the apoplasm from the BS by SWEET13 and subsequent loading into the phloem by SUT1 (Slewinski et al., 2009a; Bezrutczyk et al., 2018; Bezrutczyk et al., 2021). While these proteins are difficult to detect in the proteomic profiles, transcript levels were significantly decreased in the heat-stressed plants (Figure 8), suggesting a reduced capacity for phloem loading in heat stress. Impairments in sucrose transport in C_4_ plants have been previously linked to a dwarf phenotype (Russin et al., 1996; Ma et al., 2009a; Slewinski et al., 2009a); (Ma et al., 2009b; Slewinski et al., 2009b; Bezrutczyk et al., 2018) although, unlike our experiments, feedback inhibition of photosynthesis resulting from sugar accumulation in leaves and evidence of foliar oxidative stress were obvious in these cases.

The shift towards hexose accumulation in heat treated leaves was presumably the result of increased sucrose breakdown either by sucrose synthase or invertase. Under heat treatment, vacuolar invertase expression increased substantially at both the transcript and protein level and sucrose synthase also increased (Figure 8). This would suggest a shift toward vacuolar storage of hexoses in heat treated plants. Reduced starch accumulation did not appear to be due to reduced expression of genes encoding enzymes in starch biosynthesis (Figure 8) but likely due to allosteric regulation of the AGPase (Cross et al., 2004; Hwang et al., 2005). Activity of this enzyme is tightly controlled by the relative levels of 3-PGA (an activator of the enzyme) and Pi (an inhibitor). Under heat treatment, Pi levels increased and 3-PGA levels declined significantly (see Figure 3), which would reduce activity of this key regulatory enzyme.

The significance of the build-up of soluble sugars seen here in response to heat could in fact be in a protective and acclimation response to heat stress. Accumulation of solutes such as sugars and proline seen here is thought to provide osmo-protection of cellular processes under stress through their role as compatible solutes retaining turgor and potentially stabilising proteins (Garg et al., 2002; Lunn et al., 2014; Siddique et al., 2018). In addition to the shift from starch to sucrose and hexose, levels of raffinose rose markedly under heat treatment (Figure 7). Raffinose oligosaccharides are regarded as important molecules in stress response in plants, because of their membrane-stabilizing, antioxidant and possible predicted signalling functions (Hincha et al., 2003; Panikulangara et al., 2004; Kim et al., 2008; Nishizawa et al., 2008; Valluru and Van den Ende, 2011). The increases in raffinose oligosaccharides in our experiments are consistent with highly significant increases in expression of transcripts encoding raffinose synthase, stachyose synthase and galactinol synthase (Figures 7 and Supplementary Figure 6) to stimulate accumulation of raffinose to combat heat damage.

### Effects of hormone and sugar signaling on growth and development

Taken together, these changes in carbon partitioning indicate that *Setaria* is responding strongly to the high temperature conditions and adapting its metabolism very effectively. However, while photosynthesis appears to be protected under heat treatment, growth is certainly not. The stunting of these plants under high temperature without signs of leaf stress or photosynthetic inhibition are suggestive of a very effective homeostatic response in plant growth and minimization of vegetative damage to the plant from heat stress. A hypothesis is presented below which attempts to link metabolite levels and gene expression to sugar and hormone signaling in mediating a coordinated response to heat stress which restricts growth and minimizes damage.

Increases in ABA levels commonly observed under abiotic stress are thought to inhibit growth by stabilizing DELLA proteins which repress the growth promotive effects of GA (Szostkiewicz et al., 2010; Blanco-Touriñán et al., 2020) and act broadly across the ABA-dependent pathway of stress response (Chen et al., 2020). This ABA dependent pathway appeared to dominate the signaling response in the experiments described here, as evidenced by a 20-fold increase in ABA levels, up-regulation of gene expression of enzymes in ABA biosynthesis and increased expression of proteins known to be controlled by ABA levels (Supplementary Figure 9). Among the pathways regulated by ABA is the TOR signaling pathway (Wang et al., 2018). It has been reported that ABA accumulated following stress binds to intracellular PYR/ PYL/RCAR receptors and ABA-receptor complexes bind to and inhibit PP2C protein phosphatases (Wang et al., 2018). This releases the activity of Snf1-related protein kinase 2s (SnRK2s). The SnRK2s phosphorylate downstream targets to elicit protective responses such as stomatal closure and the expression of ABA responsive genes (Chen et al., 2020). The elevated SnRK2 activity is thought to inhibit the growth promoting action of TOR to sacrifice growth for survival during stress (Fu et al., 2020). In the current data set stomatal conductance was not substantively reduced after heat treatment (Supplementary Figure 2) which may reflect acclimation following a short-term effect of ABA. Note that expression of two of the three OST1 (OST1/SnRK2) genes in Setaria thought to be involved in ABA induced stomatal control during stress in other species (Acharya et al., 2013) decreased under heat treatment while one was increased (Supplementary Figure 9).

In addition to engagement of a strong ABA dependent growth response in *Setaria* under heat treatment, concerted action of reactive oxygen produced under high temperature stress is also known to trigger increases in JA and affect a range of growth-related targets. Increases in JA can directly affect the action of the DELLA proteins via protein stability and post-translational modification (Liu and Timko, 2021). This effect of JA would provide additional repression of growth through the GA pathway if both ABA and JA were acting to stabilise DELLA protein levels. While increases in JA were not apparent in this study, measurements of JA are challenging and strong evidence for a JA response is provided by the upregulation of gene expression for a range of JA induced transcripts (Supplementary Figure 10).

Along with the strong response of ABA targets in the transcriptional profiles after heat treatment (Supplementary Figure 9), the transcriptional response of auxin target transcripts suggested a fall in auxin levels (Supplementary Figure 11). While free IAA did not change after heats stress, there was a marked increase in amino-conjugates of auxin (Figure 9). A buildup of these conjugates has also been seen in concert with increased ABA levels in heat stressed wheat anthers (Bheemanahalli et al., 2020) and salt stressed pea seedlings (Ostrowski et al., 2016). Amino-conjugates are traditionally regarded as an inactive “reservoir” and homeostatic mechanism for regulating free IAA (Bartel, 1997) but it is also thought that these conjugates may be potent signaling compounds themselves under stress (Rosquete et al., 2013), also reducing growth and development. For example, IAA conjugate treatments strongly inhibit IAA-induced shoot growth and root initiation in tomato cell cultures (Magnus et al., 1992). There is little information in the literature on how these conjugates may function in stress signaling.

While there is strong evidence from the data shown here for a concerted downregulation of growth via hormone signalling, the large changes in carbon partitioning away from starch and toward soluble sugars (Figure 6, Figure 8 and Supplementary Figure 4) is more challenging to interpret in a signalling context. The large increases in hexose levels seen here following heat stress would be expected to feed-back on photosynthetic gene expression and trigger phenotypes similar to those seen when leaf sugar export is impeded or exogenous sugars are fed to leaves (Azcón-Bieto, 1983; Goldschmidt and Huber, 1992; Van Huylenbroeck and Debergh, 1996; Van Oosten et al., 1997; Pego et al., 2000). However, while there was some global transcriptional downregulation of photosynthetic transcripts under heat, this was often not reflected in protein levels or physiological phenotypes (Figures 2-5). There is little published evidence for sugar feedback on photosynthetic gene expression in C_4_ leaves (Lunn and Furbank, 1999; Lobo et al., 2015; Marquardt et al., 2021) with results implying that C_4_ plants are more sensitive to sugar starvation than feedback inhibition (Henry et al., 2020).

Large increases in soluble sugar levels would also be expected to have a profound effect on TOR signalling acting to promote TOR activation of ribosome biogenesis, cell division and growth (Li and Sheen, 2016; Haq et al., 2022; Jindal et al., 2022). Indeed, a highly significant (p<2.2E-16) positive correlation (r=0.44) was observed between the previously published responses of Arabidopsis TOR-regulated transcripts to a two-hour treatment with 15 mM glucose in *A. thaliana* seedlings (Xiong et al., 2013) and the responses of their *S. viridis* orthologs to prolonged heat stress in this study (Supplementary Figure 12A). Conversely, a highly significant (p<2.2E-16) *negative* association (r=-0.41) when the analogous comparison (Supplementary Figure 12B) was made with a published response of *A. thaliana* seedlings to treatment with the powerful TOR-inhibitor, AZD8055 (Dong et al., 2015). This was not evident in the response to high sugar levels in *S. viridis* grown at high irradiance (Henry et al., 2020) or inhibition of TOR action in *S. viridis* by chemical means (Da Silva et al., 2021), suggesting that TOR sensing in C_4_ plants may be less sensitive to sugar regulation than in C_3_. The negative feedback regulation of TOR signalling was also recently reported (Jindal et al., 2022), which may also explain why increased sugar levels are not always associated with TOR activation and stimulation of plant growth. It should also be noted that the majority of studies on TOR and regulation of growth and carbohydrate status have been done in sink tissue, not photosynthetic source tissue. The role of TOR in these tissues may differ substantially from that in tissues receiving externally supplied photosynthate.

### Major declines in ribosomal protein levels point to potential impairment of ribosomal biogenesis

Activation of TOR is typically associated with transcriptional upregulation of ribosome biogenesis when TOR signalling is experimentally modulated in plants (Mahfouz et al., 2006; Ren et al., 2011; Chen et al., 2018; Soprano et al., 2018; Busche et al., 2021). We observed strong coordinated transcriptional upregulation of ribosome biogenesis with the GO terms ‘translation’ and ‘ribonucleoprotein complex biogenesis’ being highly significantly over-represented among up-regulated transcripts under heat stress (Supplementary Date File 1 – Tables 3 to 5 and Supplementary Methods). In contrast, at the protein level, ribosomal proteins tended to be decreased in abundance with 83 (69%) of the 121 quantified ‘ribosomal protein’-annotated proteins being decreased in abundance by between 20 and 90% and only 6 (5%) being increased in abundance by more than 20% (Supplementary Data File 1 - Table 2). Prolonged (37°C) heat stress has recently been shown to cause impairment of ribosome biogenesis in Arabidopsis by compromising the processing of ribosomal RNA (rRNA) precursors in the nucleolus (Darriere et al., 2022). A similar impairment of ribosome biogenesis in *S. viridis* may contribute to the major decline in ribosomal protein levels observed here. Such an impairment would be expected to be detrimental to growth by decreasing the energy efficiency of ribosome biogenesis and growth and may contribute to the impaired growth phenotype observed in *S. viridis* under prolonged heat stress in this study.

Almost all of the observed increases in ribosomal protein abundance were marginal (up to 1.6-fold). Interestingly, however, the abundance of one particular ribosomal protein, RPL10A (encoded by Si026830m), increased by over 20-fold (Supplementary Data File - Table 2). It has recently been shown that RPL10A expression is upregulated by ABA and that RPL10A may be a positive effector of ABA-mediated responses in Arabidopsis (Ramos et al., 2020). It would be interesting to explore the role of ABA-mediated RPL10A accumulation in the *S. viridis* heat stress response via mutant studies. It remains unclear from our results whether decreased ribosome abundance is a negative consequence of thermal disruption to ribosome biogenesis/stability or a survival-promoting response orchestrated by complex homeostatic sensing and signalling networks involving TOR and ABA. Either way, it is clear that the reduced plant growth observed under heat stress in this study may be explained at least in part by reduced protein synthesis capacity.

## Conclusion

Long term high temperature stress of *S. viridis* induces a strong “dwarf” phenotype which is a result of a converging set of hormonal signals all impacting to reduce growth. Remarkably, photosynthetic processes were quite robust under high temperature and appear to have been protected from damage by acclimation of carbohydrate metabolism, oxidative stress responses and potentially by flexibility in the production of aspartate versus malate in the C_4_ photosynthetic pathway in this species under conditions where the mesophyll chloroplast stroma was experiencing oxidizing conditions.

## Acknowledgements

The study was funded by the ARC Centre of Excellence for Translational Photosynthesis (CE140100015). RES is supported by the VC Fellowship at Western Sydney University. Hormone analysis was completed by Dr. Thy Truong at the Australian National University Research School of Biology / Research School of Chemistry Joint Mass Spectrometry Facility as an in-house service and the training and support of the ANU Research School of Biology / Research School of Chemistry Joint Mass Spectrometry Facility (JMSF) in carrying out metabolomics analyses.

## Author Contributions

The study was conceived by RTF, SvC and RES. PZ, AC and RES performed experiments. All authors contributed to data analysis, writing and editing of the manuscript.

